# Accurate and Efficient Gene Function Prediction using a Multi-Bacterial Network

**DOI:** 10.1101/646687

**Authors:** Jeffrey Law, Shiv Kale, T. M. Murali

## Abstract

**Motivation:** Nearly 40% of the genes in sequenced genomes have no experimentally- or computationally-derived functional annotations. To fill this gap, we seek to develop methods for network-based gene function prediction that can integrate heterogeneous data for multiple species with experimentally-based functional annotations and systematically transfer them to newly-sequenced organisms on a genomewide scale. However, the large size of such networks pose a challenge for the scalability of current methods.

**Results:** We develop a label propagation algorithm called FastSinkSource. By formally bounding its the rate of progress, we decrease the running time by a factor of 100 without sacrificing accuracy. We systematically evaluate many approaches to construct multi-species bacterial networks and apply FastSinkSource and other state-of-the-art methods to these networks. We find that the most accurate and efficient approach is to pre-compute annotation scores for species with experimental annotations, and then to transfer them to other organisms. In this manner, FastSinkSource runs in under three minutes for 200 bacterial species.

**Availability and Implementation:** Python implementations of each algorithm and all data used in this research are available at http://bioinformatics.cs.vt.edu/~jeffl/supplements/2020-fastsinksource.

**Supplementary Information:** A supplementary file is available at **bioRxiv** online.

## 1 Introduction

The number of fully sequenced prokaryotic genomes is increasing dramatically (Land *et al.*, 2015). The overwhelming majority of the genes in these genomes have not been studied experimentally. In fact, fewer than 0.01% (about 10,000 out of 104 million) of prokaryotic genes in the UniProt Knowledgebase (UniProt Consortium, 2018) have a Gene Ontology (GO) (Gene Ontology Consortium, 2018) annotation with an experimental evidence code. To address this gap, several computational methods have been developed to associate molecular functions and biological processes with genes lacking experimental annotations. The GO consortium uses many approaches including homology inference (Finn *et al.*, 2016) and phylogenetic analysis (Gaudet *et al.*, 2011) to annotate genes to GO terms using computational or electronic evidence codes.

Despite the sophistication of these methods, nearly 40% of all genes in sequenced genomes have no annotation at all (Cozzetto and Jones, 2017).

To fill this gap, a broad range of techniques have been developed that integrate multiple types of data (e.g., physical interactions, co-expression, and cellular pathways) (Mostafavi *et al.*, 2008; Wang *et al.*, 2015; Cho *et al.*, 2016; Gligorijević *et al.*, 2018). These methods can predict gene function on a genomewide scale but usually operate on a single organism. They are also limited by the lack of availability of diverse, rich functional genomic datasets for all but a handful of organisms.

Our main goal in this paper is to develop highly-efficient algorithms that can make genomewide functional predictions for a newly-sequenced “target” species by effectively using heterogeneous experimental data from well-studied organisms. We adopt a network-based strategy since it facilitates the representation and integration of diverse datasets and because these types of approaches have been effective in gene function prediction (Mostafavi *et al.*, 2008; Cho *et al.*, 2016; Jiang *et al.*, 2017).

Two distinct challenges arise at this point. The first is that we have a large number of options for constructing a network that involves the genes in the target and well-annotated species depending on which type of information we want to integrate. The second challenge is that the largest networks we build can contain tens of millions of edges. This size requires efficient algorithms for network-based gene function prediction.

To address the first challenge, we systematically explore many combinations of networks for connecting the genes in well-annotated organisms to each other and to those in the target species (Section 2.4). We evaluate these strategies using a leave-one-species-out (LOSO) validation framework that mimics the challenge of making functional predictions for a newly-sequenced species where none of its genes have any annotations. Initially, we apply LOSO evaluation to 25 bacteria with a sufficient number of experimental annotations, which we call the “core” species. To test scalability and assess the trade-off between speed and accuracy of these approaches, we then expand to 200 species. We find that it is not necessary consider the genes in target species during network propagation, i.e., we were able to generate accurate functional predictions by running the algorithms for only the core species, and then transferring those predictions to the target organism(s) using sequence similarity.

To address the second challenge, we developed a novel, iterative label propagation algorithm called FastSinkSource for which we mathematically prove that the rate of convergence of the computed score for every node in the network is geometric in the number of iterations (Section 2.1). This property enabled us to decrease the running time of FastSinkSource by a factor of 100 or more without sacrificing prediction accuracy.

We found that FastSinkSource and two other state-of-the-art network-based algorithms, BirgRank and GeneMANIA computed far more accurate predictions than a baseline BLAST-based method. In addition, FastSinkSource was much more efficient than BirgRank with similar or better prediction accuracy. FastSinkSource was moderately more accurate than GeneMANIA while achieving the same runtime efficiency.

## 2 Methods

### 2.1 FastSinkSource

A standard formulation of network-based GO annotation prediction is as a semi-supervised problem. We are given a weighted undirected network *G* = (*V, E, w*), where *w_uv_* is the weight of the edge (*u, v*). In addition, for a GO term *τ*, we have a partition of the nodes in *V* into three sets: *V*^+^, *V*^−^ and *V*^0^ positive, negative, and unlabeled examples, respectively. The goal is to compute a score *s*(*u*) for each node *u* that is an unlabeled example, by propagating the positive and negative labels across *G*. Strictly speaking, the score depends on *τ* as well but we omit it for clarity.

Given a parameter 0 < *α* ≤ 1, FastSinkSource computes a score *s*(*u*) between 0 and 1 for each node *u* that is an unlabeled example for *τ* by minimizing the function

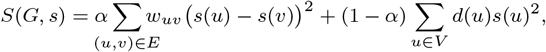

with the scores of positive and negative examples fixed at 1 and 0, respectively. Note that if *α* = 1, FastSinkSource is identical to the SinkSource algorithm (Zhu *et al.*, 2003; Murali *et al.*, 2011).

We define **P** to be a |*V*^0^|×|*V*^0^| matrix of the degree-normalized edge weights in *G* among pairs of nodes in *V*^0^, i.e., *P_uv_* = *w_uv_/d*(*u*) for every pair of unlabeled examples (*u, v*). We also define **s** to be a |*V*^0^| × 1 vector formed by the node scores of all unlabeled examples and **f** to be a |*V*^0^| × 1 constant vector in which *u*th element *f*(*u*) is the contribution to *s*(*u*) from the positively-labeled neighbors of *u*. Specifically, 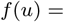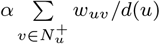. Then **s** satisfies the following linear system:

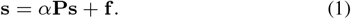

We can compute the solution **s** = (**I**−*α***P**)^−1^**f** using power iteration (Zhu *et al.*, 2003). We now describe how to compute an upper and lower bound on every node’s score after each iteration and the benefits of these bounds.

#### Lower and upper bounds

If *s*(*u, i*) denotes the score of node *u* after *i* iterations, then we can prove that the score for every node increases with the number of iterations, i.e., *s*(*u, i*) ≤ *s*(*u, i* + 1), and that

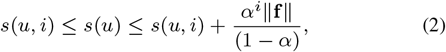

where ‖**f**‖ is the largest absolute value of the entries in **f**. In other words, the score *s*(*u, i*) is a lower bound on the final score *s*(*u*) and 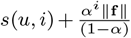 bound *s*(*u*) from above. Thus the difference between the sought-for score and its value after *i* iterations decreases geometrically with *i*. We based our proof of this result (Section S1) on Zhang *et al.* (2015), who used similar ideas for finding the *k* nodes with the highest RWR scores.

Now consider the scores of two nodes *u* and *v* after *i* iterations (Figure 1). If 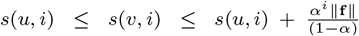, then the interval spanned by the lower and upper bounds for *s*(*u*) overlaps the corresponding interval for *s*(*v*). In this case, it is unclear if the final score *s*(*u*) will be larger or smaller than *s*(*v*). However, if we observe in a later iteration *j* that 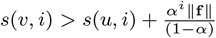, then the intervals are disjoint, guaranteeing that *s*(*u*) < *s*(*v*) (Figure 1). We can conclude that if the interval spanning the lower and upper bounds of *u* does not overlap with any other node’s interval after a certain number of iterations, then we have determined *u*’s rank upon convergence. If this property is true for every node, then we have ranked all the nodes correctly and no further iterations are needed. This observation has two significant benefits:

a. It inspires an alternative strategy for checking convergence. At the end of each iteration, we sort all the nodes in increasing order of score. Then for each index *k* and node *u_k_* in this sorted order, we check if 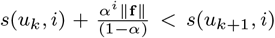. If this inequality is true for every index, then the rankings will not change in subsequent iterations and we say that the process has converged. Note that we need only compare *O*(|*V*|) pairs of node scores to check for convergence. This approach is especially useful when we only need to compute the correct ranking of a subset of nodes by their scores, e.g., during LOSO validation.
b. If we stop after *i* iterations, we have an estimate 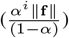 of how much we have under-estimated every node’s score. We also have an upper bound on the number of node pairs that may be ranked incorrectly with respect to each other, which is the number of node pairs that have overlapping lower and upper bounds.

**Fig. 1.**
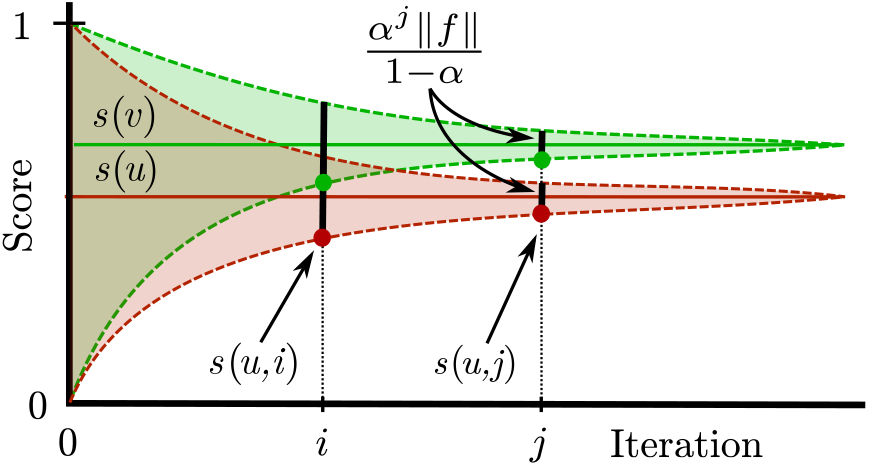
Illustration of the geometric convergence of scores in FastSinkSource. The solid red line indicates the score *s*(*u*) for a node *u* that we are seeking to compute. The shaded red region shows how different *s*(*u*) can be from the score computed for node *u* after each iteration. The green line and region correspond to node *v*. At iteration *i*, we cannot be certain if *s*(*v*) is larger or smaller than *s*(*u*) because the difference between their computed scores after *i* iterations is less than the error term. By iteration *j*, however, the difference is larger than the error term, which implies that *s*(*v*) > *s*(*u*).

It is possible for two or more nodes to have the same lower bound and the same upper bound in every iteration. Our approach can determine the rank of these nodes as a group with respect to the other nodes. In this case, we simply give these matching nodes the same ranking as we have no way to distinguish how to order them in relation to each other.

### 2.2 Other Algorithms

We compared FastSinkSource to its predecessor SinkSource (Murali *et al.*, 2011), two other network propagation methods: GeneMANIA (Mostafavi *et al.*, 2008), and BirgRank (Jiang *et al.*, 2017), and a baseline method we call Local. GeneMANIA utilizes Gaussian Random Field label propagation to diffuse labels from positive and negative examples to make term-based predictions. SinkSource is similar to GeneMANIA but does not allow the scores of the given positive and negative examples to change. BirgRank (BI-Relational Graph page RANK) constructs a bi-relational graph with a given network and a GO hierarchy, connecting each gene to every GO term for which it has an annotation. It applies Random Walk with Restarts (RWR) (Page *et al.*, 1999) to diffuse the annotation information across this network. Local mimics a BLAST-based procedure for transferring GO term annotations, where a given gene’s score for a term is the weighted average of the scores of its neighbors. See Section S2 for more details about the algorithms and our implementations, respectively.

### 2.3 Datasets

#### Gene Ontology annotations

We considered three sets of evidence codes: (i) Experimental as well as those used in the CAFA evaluations (EXPC) (Jiang *et al.*, 2016): EXP, IDA, IPI, IMP, IGI, IEP, TAS, IC; (ii) Computational Analysis (COMP): ISS, ISO, ISA, ISM, IGC, IBA, IBD, IKR, IRD, RCA; and (iii) Electronic Analysis (ELEC): IEA. We selected the 200 bacterial species with the largest number of GO annotations (protein-GO term pairs) with EXPC or COMP evidence codes in the UniProt-GOA database (“goa_uniprot_gcrp.gaf.gz” downloaded on October 15, 2019). These 200 species did not contain any annotations with the recently-introduced high-throughput evidence codes. See Section S3.1 for details of our definition of positive and negative examples.

#### Bacterial proteomes

We selected the 200 bacterial reference proteomes with the most EXPC and COMP annotations and obtained their protein sequences from UniProt (downloaded on October 15, 2019). We split these organisms into two groups: i) the *core* species, i.e., those with at least one GO term with 10 or more annotations with EXPC evidence codes, and ii) the non-core species. We define a *target* species as one or more organisms in the non-core group for which we wish to make predictions. There were 25 and 40 species, respectively, in the core when we considered BP and MF terms (Tables S1 and S2). About 91% and 93% of all bacterial BP and MF EXPC annotations, respectively, came from the set of core species.

#### Sequence similarity network

We ran all-vs-all BLASTP with default parameters, except for the E-value cutoff set to 20 and the parameter `-max_hsps` (i.e., maximum High Scoring Segment Pairs), set to 1. For the database required by BLAST, we used the protein sequences of the 200 species. We processed the results by retaining the weaker score for all reciprocated matches, removing self-comparisons, and using the negative of the base-10 logarithm of the E-value as the edge weight. If the E-value was 0, we assigned a weight of 180, which was the largest (rounded) value we observed. We tested various E-value cutoffs from 1 × 10^−25^ to 20. If the cutoff was larger than one, we added the base-10 logarithm of the cutoff to every edge weight to make it positive.

#### STRING networks

We integrated species-specific functional association networks with the sequence similarity network (SSN) for bacterial species that had networks available in the STRING database (v11, downloaded on May 30, 2019) (Szklarczyk *et al.*, 2019). A STRING network was available for 23 of the 25 core species (for BP terms) and 173 of the 200 species. We converted the STRING IDs to UniProt IDs using the mappings available from UniProt. STRING assigns edge weights based on multiple lines of evidence of association including physical binding, gene expression, and orthology mapping. We utilized six STRING networks: neighborhood, fusion, cooccurence, coexpression, experimental, and database. We also tested various cutoffs of the STRING combined edge scores including 150, 400, 700, and 900 (low, medium, high and very high stringency).

#### Network integration

We tested two methods to integrate the SSN and STRING networks: term-by-term weighting described in the original GeneMANIA publication (GMW; we use this abbreviation to differentiate the weighting method from the network propagation algorithm) (Mostafavi *et al.*, 2008) and Simultaneous Weights with Specific Negatives (SWSN) (Youngs *et al.*, 2013) which weights networks using multiple related terms simultaneously (Section S3.2).

### 2.4 Network Combinations

We faced several choices in constructing the network linking the genes of the target species to each other and to genes in the core group. Does the target species have a STRING network? Should we connect the genes in the core species based on sequence similarity or STRING edges or both? How should we treat species that have no EXPC annotations? To study the effects of these choices on the accuracy and scalability of gene function prediction algorithms, we proceeded as follows.

We divided the sequence-similarity network (SSN) and STRING networks into the following subsets (Figure 2): (a) *SSN-T* : The sequence-similarity-based edges between the genes in the target species and those in the core species. (b) *SSN-C*: The sequence-similarity-based edges among the genes in the core species. (c) *STRING-C*: The union of the STRING networks for core species. (d) *STRING-T* : The STRING network for the target species. This demarcation of networks generalizes to the case when there are multiple target species for which we want to make predictions simultaneously. In addition, we observed that the core group remains the same even as we vary the target species. Therefore, we consider an approach where we pre-compute prediction scores for the genes in the core species, and then transfer them to the target genes using sequence similarity. We refer to this approach as *SSN-Nbrs* (i.e., SSN-Neighbors). In this case, we first computed scores using only nodes and edges in the core, then for each gene in the target species, we took the weighted average of the scores of its neighbors using the SSN-T edges.

**Fig. 2.**
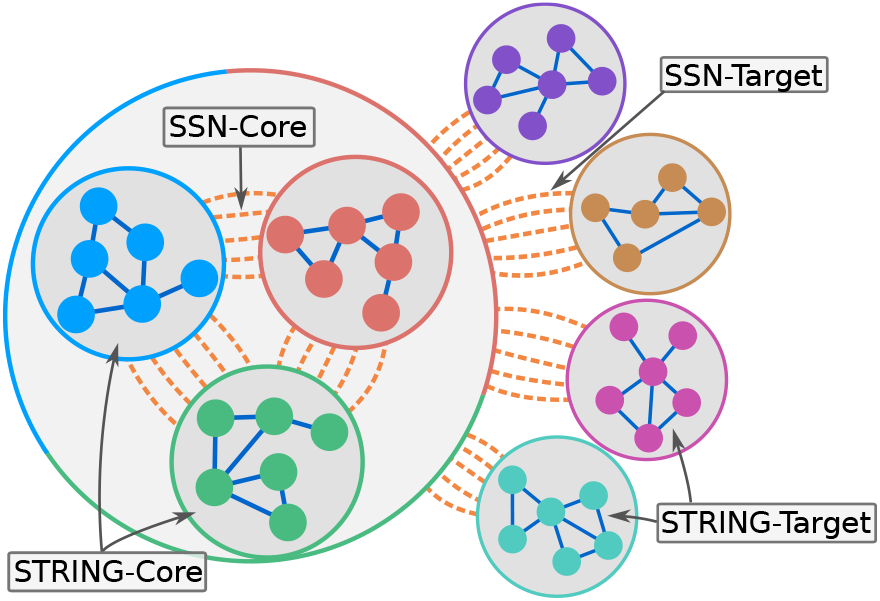
Illustration of network-based approach to predict functions for an arbitrary number of “target” species. Dashed and solid edges represent inter- and intra-species edge types, respectively (e.g., SSN and STRING). Edges between groups of nodes represent edges between the nodes in the respective groups.

Based on this partition, we evaluated the following nine network combinations, where ‘+’ denotes the union of networks, and ’,’ denotes the sequential use of networks:

i. SSN-T: sequence-similarity edges between genes in the target species and those in the core species.
ii. SSN-T+SSN-C: in addition to (i), the sequence-similarity edges among the genes in the core.
iii. SSN-T+STRING-C: in addition to (i), the STRING networks for every core organism.
iv. SSN-T+SSN-C+STRING-C: in addition to (ii), the STRING networks for every core organism.
v. SSN-T+SSN-C+STRING-C+STRING-T: in addition to (iv), the STRING network in the target species. This network contains all the edges we considered, which we refer to as the full network.
vi. SSN-T+STRING-C+STRING-T: the network in (v) without the sequence-similarity edges among the genes in the core species.
vii. SSN-T+STRING-T: the network in (vi) without the STRING networks in the core species.
viii. STRING-C,SSN-Nbrs: the STRING networks for every core organism followed by the network in (i).
ix. SSN-C+STRING-C,SSN-Nbrs: in addition to (viii), the sequence-similarity edges among the genes in the core species.

Evaluating these choices in sequence, we expected to dissect the relative utility of using SSN-T compared to SSN-Nbrs, as well as the SSN-C, STRING-C, and STRING-T networks. We did not use the SSN-T+SSN-C+STRING-T network since it was very unlikely that a STRING network would exist for the target species but not a core species. We also did not use SSN-C,SSN-Nbrs to limit the number of combinations. Note that we either needed to include the SSN-T network or use the SSN-Nbrs approach. Otherwise, we would not be able to make predictions for a target species.

### 2.5 Evaluation

We split our evaluations into two types as follows:

i. *Leave-One-Species-Out (LOSO) validation.* In this method, for each species *s* of the core group, we leave out the annotations of all genes of *s*, apply each algorithm using the annotations of the other core species, and then assess how well we can recover the annotations of *s*. In effect, *s* is removed from the core group and serves as a target species during this evaluation. We perform this evaluation only for EXPC annotations.
ii. *Target species validation.* Experimentally-based GO term annotations to evaluate the predictions of the algorithms are sparse for non-core species. Therefore, we use annotations with curator-reviewed computational analysis (COMP) or electronic (ELEC) evidence codes as alternative ground-truth datasets. In this validation, we apply every algorithm using only the EXPC annotations for the core species and assess how well we can recover the COMP or ELEC annotations in the target species. Here we considered only those target species (i.e., in the non-core group) for which at least one term had 10 or more annotations of the given evidence code type. For the BP COMP, MF COMP, BP ELEC, and MF ELEC evaluations, there were 31, 24, 175, and 160 target species, respectively.

#### Term selection

For a term *t* to be evaluated, we required at least 10 EXPC annotations to *t* in the core species (minus the left-out species in the case of LOSO validation), and at least 10 annotations to *t* of the given evidence code group in the target organism. For each species, we restricted our evaluation to its *leaf terms*, which we defined as those that did not have any children in the GO DAG that also satisfied our criteria.

#### Evaluation

When evaluating the predictions for a given species, we calculated true positives and false positives from the positive and negative examples of the given evidence codes for that species. We summarized the quality of the predictions for each GO term using the *F*_max_ value, which is the maximum of the harmonic mean of the precision and recall (i.e., F_1_ score) over the entire precision-recall curve.

## 3 Results

We evaluate the performance of four network propagation algorithms and a baseline method on nine different combinations of networks (Section 2.4) for different subsets of annotations: LOSO validation for EXPC annotations in core species, COMP annotations in target species (not in the core), and ELEC annotations in target species. We also compared the running times of the algorithms. We focus on the results for BP GO terms, and briefly mention our findings for Molecular Function (MF) terms.

### 3.1 Parameter Selection

We first determined values for the E-value cutoff for the SSN, the weight threshold for STRING networks, the method for network integration, and the different parameters required by FastSinkSource and BirgRank. We used EXPC LOSO validation of BP terms on the full network (i.e., SSN-T+SSN-C+STRING-C+STRING-T) for the selection of all parameters, except for the E-value cutoff, for which we used SSN-T+SSN-C.

#### E-value cutoff

To evaluate the potentially deleterious effects of poor homology on prediction quality, we used different E-value cutoffs to construct the SSN; Table S3 shows the sizes of these networks. We found that the median *F*_max_ of every method gradually increased as we raised the E-value cutoff, but did not improve past 0.1 (Section S4.1). Thus, we selected a cutoff of 0.1.

#### Network integration

After evaluation of GMW and SWSN across multiple cutoffs of STRING edge confidence, we selected SWSN with a cutoff of 700 to integrate the networks (Section S4.2), which resulted in a total of 96,107 nodes and 1,596,336 edges (Table S3). Note that during LOSO validation, for each species we left out, we weighted the networks using only annotations of the organisms that we retained.

#### FastSinkSource and BirgRank

We tested eight different values of *α* and found *α* = 0.99 to yield the highest median *F*_max_ value. We next varied the number of iterations from 400 to 1 and found that any value below 20 resulted in a statistically significant decrease in the *F*_max_ values when compared to *α* = 0.99 and 400 iterations or to *α* = 1 and 1,000 iterations. Therefore, we chose *α* = 0.99 and 20 iterations for subsequent analyses. See Section S4.3 for details. We performed parameter estimation for BirgRank as well (Section S4.4).

### 3.2 Evaluation of Network Combinations

#### LOSO validation for EXPC annotations

We first performed LOSO validation for the 25 core species with EXPC annotations. The median *F*_max_ of Local was 0.43 (Figure 3(a)). In contrast, other than for SSN-T and STRING-C,SSN-Nbrs, every network propagation algorithm achieved a median *F*_max_ of 0.5 or more, with the largest median value exceeding 0.65. Moreover, network propagation methods also improved when provided additional networks. The full network (SSN-T+SSN-C+STRING-C+STRING-T) yielded the highest median *F*_max_ values, except for SinkSource, which achieved a slightly higher median *F*_max_ of 0.67 for SSN-T+SSN-C+STRING-C.

**Fig. 3.**
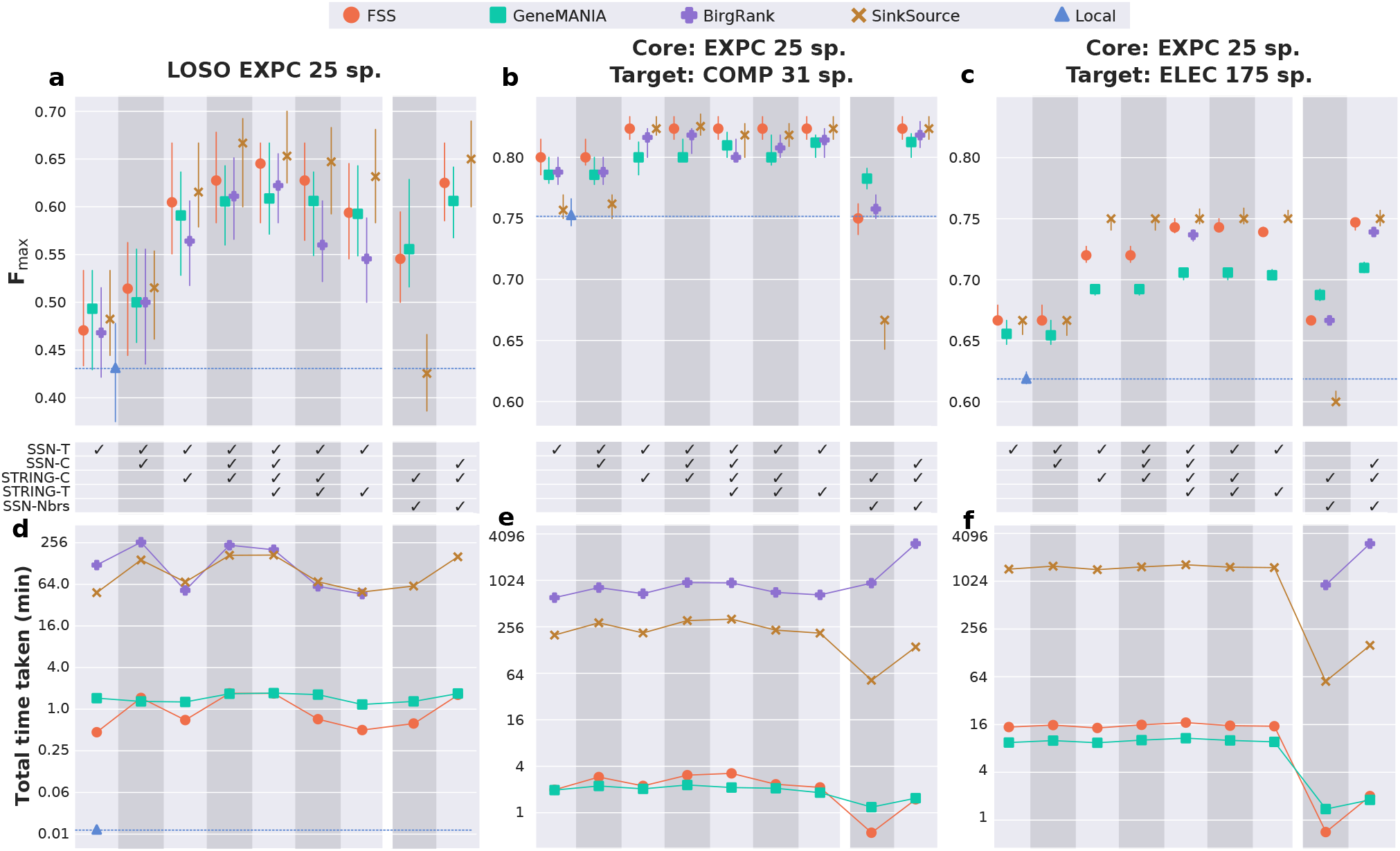
Comparison of *F*_max_ results for BP EXPC LOSO evaluation, as well as the evaluation of species with only COMP and ELEC annotations across five algorithms and nine networks. (**a-c**) Median *F*_max_ as well as the 95% confidence interval of the median, estimated using 1,000 bootstrapped samples of the data. Boxplots of the full *F*_max_ distributions appear in Figure S1. (**d-f**) Running time for each algorithm, measured in independent runs using a single core on a computer with a 2.2 GHz processor and 64GB of memory running CentOS version 7. (**a**) Results for EXPC LOSO evaluation; 25 species and 242 BP terms (355 total pairs). (**b**) Results for evaluation of species with COMP and no EXPC; 31 species and 301 BP terms (1,378 total pairs). (**c**) Results for evaluation of species with ELEC and no EXPC; 175 species and 339 BP terms (15,237 total pairs). (**d**) Total running time of each method for EXPC LOSO (**a**). (**e,f**) Running time of each method to compute scores for the target-only species (**b,c**). We limited algorithms to a maximum of 72 hours of running time, which explains the missing points for BirgRank.

To study these trends further, for each network propagation algorithm, we determined if there were statistically significant differences between the *F*_max_ distributions for each network combination. A *p*-value < 0.05 (6.1 × 10^−9^ for FastSinkSource, 2.1 × 10^−5^ for GeneMANIA, 2.6 × 10^−5^ for BirgRank, and 2.8 × 10^−25^ for SinkSource) for the Kruskal Wallis test, after correction by the number of algorithms, indicated that these nine sets of *F*_max_ values did not originate from the same distribution. Therefore, we performed post-hoc pairwise comparisons using a Benjamini-Hochberg corrected one-sided Wilcoxon signed-rank test, and found that for FastSinkSource and GeneMANIA, the seven network combinations that included STRING, except for STRING-C,SSN-Nbrs, significantly outperformed those without STRING, namely SSN-T and SSN-T+SSN-C (*q*-value < 0.05, Table S5). For the 15 pairs of networks in this group of six combinations, we did not observe a statistically-significant difference for 13 pairs. The exceptions were the improvement of the full network (fifth column in Figure 3(a)) over the SSN-T+STRING-C and SSN-T+STRING-T networks (*q*-values of 0.03 and 0.01 for FastSinkSource, and 0.005 and 0.01 for GeneMANIA, respectively). For SinkSource the trends were largely similar, except that only the decrease of *F*_max_ for the SSN-T+STRING-C combination in comparison to the full network was statistically significant. For BirgRank, the two largest combinations, SSN-T+SSN-C+STRING-C and SSN-T+SSN-C+STRING-C+STRING-T, had a statistically significant improvement in *F*_max_ compared to all other combinations (*q*-value < 0.05). From these comparisons, we concluded that a STRING network for the target species was not needed as long as the SSN and STRING networks are included among the core species. This finding is especially relevant since a STRING network or other network datasets are likely not to be available for a newly sequenced target species.

#### Target species validation for COMP annotations

Next, we evaluated the ability of our methods to recover BP annotations with COMP evidence codes in 31 target bacterial species when using the EXPC annotations in the 25 core species to compute prediction scores. The median *F*_max_ for most methods and network combinations improved to over 0.8. The median increase in *F*_max_ for any network propagation algorithm over the Local baseline did not exceed 0.1 (Figure 3(b)). We reasoned that the relatively high median *F*_max_ of Local could be attributed to the fact that over 99% of the 113,360 curator-reviewed annotations based on computational analysis for these organisms had the evidence codes IBA (Inferred from biological aspect of ancestor, 99,563 annotations) or ISS (Inferred from sequence or structural similarity, 12,946 annotations). These annotations are likely derived from genes with EXPC annotations in the core species.

For each algorithm, we compared the *F*_max_ distributions across the network combinations just as we did for EXPC annotations above. Every algorithm had a Bonferroni-corrected *p*-value < 10^−4^ for the Kruskal-Wallis test. Therefore, we performed post-hoc tests for every pair of network combinations. In almost all cases, only the differences between the first two columns (SSN-T and SSN-T+SSN-C) and six of the other combinations were statistically significant (Benjamini-Hochberg corrected one-sided Wilcoxon signed-rank test *q*-value < 0.05). The exception was STRING-C,SSN-Nbrs which performed significantly worse than all other combinations for three of the four algorithms, especially for SinkSource. In contrast, the SSN-C+STRING-C,SSN-Nbrs approach had *F*_max_ distributions that were statistically indistinguishable from the combinations in columns three to seven, suggesting that pre-computing network propagation scores for core species is an effective strategy.

#### Target species validation for ELEC annotations

We then turned our attention to recovering ELEC annotations in 175 target bacterial species using EXPC annotations in the 25 core species. We observed that the median *F*_max_ values, by and large, were in-between the previous two evaluations, with Local at 0.62 (Figure 3(c)). The network propagation methods achieved median *F*_max_ improvements of up to 0.13 over Local. Notably, the combination of networks which resulted in the highest median *F*_max_ for three of the four methods was SSN-C+STRING-C,SSN-Nbrs. Recall that in this method, we pre-compute scores for the genes in the core species and transfer the scores to the genes in the target species by taking the weighted average of neighbors in the SSN. Using either SinkSource or FastSinkSource for pre-computing scores for the core species yielded a median *F*_max_ of 0.75, an improvement of 0.05 over GeneMANIA. BirgRank also performed the best on this combination (median *F*_max_ of 0.74). We did not test for statistical significance of the differences here due to the large number of species-term pairs being evaluated (over 15,000). In the next section (Section 3.3), we compare the prediction quality of each method on the SSN-C+STRING-C,SSN-Nbrs combination in more detail.

#### Running times

We compared the running times for every algorithm on each network combination (Figure 3(d,e,f)). For EXPC LOSO, we report the total running time of each method to compute the prediction scores (Figure 3(d)). FastSinkSource achieved the fastest running time of the propagation methods, ranging from 0.46 minutes on SSN-T+STRING-T (the second smallest network) to 1.7 minutes on SSN-T+SSN-C+STRING-C+STRING-T (the largest network), with a factor of speed improvement ranging from 75 to 262 over BirgRank, and 97 to 104 over SinkSource, respectively. GeneMANIA was more efficient than FastSinkSource in some cases, with the largest speed improvement of a factor of 3.1 on the SSN-T network. Compared to Local, FastSinkSource was 21 to 28 times slower. Note that FastSinkSource, SinkSource, and GeneMANIA performed network-wide predictions, i.e., they computed a score for each unknown example in every species. In contrast, for BirgRank, we limited predictions to the set of proteins that were either positive or negative examples in the target species, except in the case of SSN-Nbrs where scores were computed for all nodes in the core. See Section S5 for an explanation of the prohibitive running time for the missing points.

For COMP and ELEC validation, we compute scores for the genes in all the target species simultaneously. We report these running times in Figure 3(e,f). Not surprisingly, we found that the combinations which pre-computed the scores for the core species and then transferred these scores using SSN-Nbrs were faster in general than those which directly included the nodes of the target species into the network propagation (i.e., combinations that include SSN-T). This difference was especially apparent when computing scores for 175 target species, i.e., ELEC validation (compare last two columns of Figure 3(f) to the first seven columns).

The number of species increased by a factor of 5.4 when we moved from COMP to ELEC annotations with concomitantly larger networks (4 times more nodes and 6 times more edges on average, Table S4). SinkSource, FastSinkSource, and GeneMANIA took 4–8 times longer to run on these larger networks (compare first seven columns of Figure 3(e,f)). In contrast, for the SSN-Nbrs-based methods (last two columns in Figure 3(e,f)), the running time for every algorithm remained about the same. FastSinkSource and GeneMANIA took a mere two minutes using the SSN-C+STRING-C,SSN-Nbrs combination (Figure 3(e,f), last column). Note that we did not run BirgRank to completion on most combinations for the ELEC evaluation due to prohibitive running times. We made an exception for the full network, for which BirgRank took over 300 hours, to provide a reference point of the likely upper limit of its running time.

#### Results for MF terms

We repeated these evaluations for MF terms. For the EXPC LOSO evaluation, using the SSN-T+SSN-C network yielded median *F*_max_ values of 0.74 or higher for the network propagation methods, with improvements up to 0.14 over Local (median *F*_max_ of 0.66 Figure S2(a)). The integration of the STRING networks did not increase the median *F*_max_ above that of the SSN-T+SSN-C combination for three out of four methods for the EXPC LOSO evaluation, suggesting the data in STRING were not helpful for predicting MF annotations. When recovering COMP and ELEC annotations, the network propagation methods improved over Local by up to 0.07 and 0.13, respectively (Local median *F*_max_ of 0.8 and 0.76, respectively; Figure S2(b,c)). Integrating the STRING networks gave only marginal improvements, with a median *F*_max_ increase of up to 0.02 for COMP, and up to 0.03 for ELEC, respectively. Although we did not test the SSN-C,SSN-Nbrs combination, this would likely be the best approach for computing prediction scores of MF terms for a large number of target species.

### 3.3 Examination of SSN-Nbrs Approach for ELEC

In this section, we examine the results for the SSN-C+STRING-C,SSN-Nbrs approach in the ELEC evaluation in more detail. We selected this network combination since it both accurate and efficient, as shown in the previous section. We first investigated the improvement of FastSinkSource over GeneMANIA. For almost all of the 175 target species evaluated, the median *F*_max_ (taken over the BP terms we considered) was higher for FastSinkSource than GeneMANIA, with a median improvement in the median *F*_max_ of 0.05 or higher for 17% of species (Figure 4a). When comparing the individual species-term pairs, we found that FastSinkSource improved over GeneMANIA (i.e., higher *F*_max_ values) 62% of the time, whereas the reverse was true for only 22% of the pairs.

**Fig. 4.**
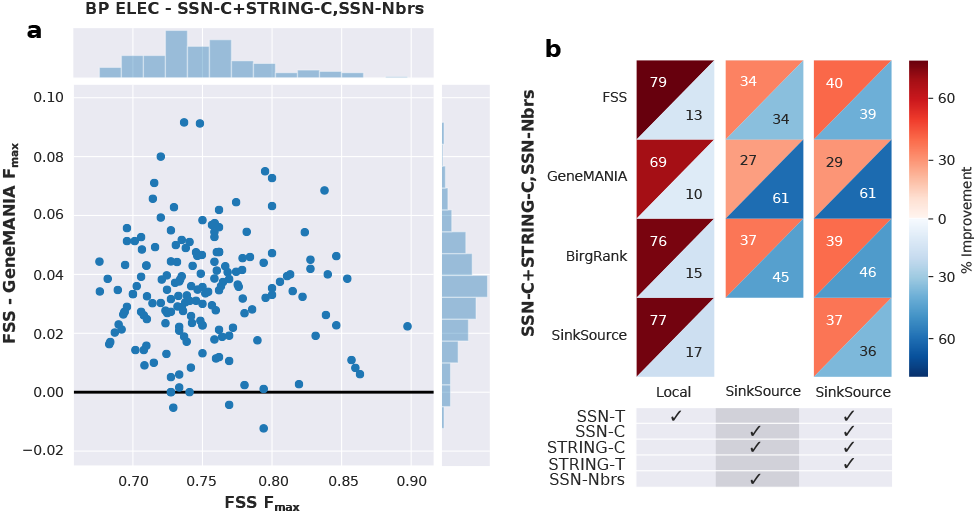
Comparison of *F*_max_ distributions for the SSN-C+STRING-C,SSN-Nbrs network combination for the ELEC evaluation for BP terms. (**a**) Scatterplot of the difference between the median *F*_max_ of FastSinkSource and GeneMANIA vs. the median *F*_max_ for FastSinkSource. Each point represents one species. (**b**) Table comparing methods using the SSN-C+STRING-C,SSN-Nbrs approach (rows) to Local and SinkSource on the same network combination, and to SinkSource on the full network (columns). Each cell contains the percentage of species-term pair *F*_max_ values that increase (upper-left triangle) or decrease (lower-right triangle) when comparing a row to a column.

Next, we compared each of the network propagation algorithms using the SSN-C+STRING-C,SSN-Nbrs approach with (a) the baseline Local on the same network, (b) SinkSource on the same network (since this algorithm achieved the highest *F*_max_ for that combination), and (c) the method and combination which resulted in the highest median *F*_max_ over all ELEC evaluations (i.e., SinkSource on the full network). For the SSN-C+STRING-C,SSN-Nbrs combination, every network propagation method substantially improved over Local, with FastSinkSource achieving a higher *F*_max_ for as many 79% of species-term pairs (first column of Figure 4b). For the same combination, FastSinkSource and SinkSource were evenly matched, with each method surpassing the other for 34% of species-term pairs (second column of Figure 4b). The trends were similar when comparing FastSinkSource on the SSN-C+STRING-C,SSN-Nbrs combination with SinkSource on the full network (third column of Figure 4b). SinkSource achieved larger *F*_max_ values for a greater percentage of species-term pairs than BirgRank and GeneMANIA (second and third columns of Figure 4b). These results demonstrate that FastSinkSource outperforms GeneMANIA in terms of prediction quality without sacrificing accuracy compared to SinkSource.

## 4 Discussion

We have systematically evaluated many network combinations for large-scale, multi-species gene function prediction. We have also developed a highly-scalable algorithm called FastSinkSource for this task. We found that the most accurate and scalable method is to use FastSinkSource to pre-compute prediction scores for the genes of core species (i.e., with experimentally validated functions) connected in a network built from many molecular systems datasets, and then to transfer those scores to the genes of a target species using sequence homology. The benefits of our approach include the ability to rapidly compute scores for any number of target species, and increasing the number of positive training examples by pooling information in multiple organisms.

Our inability to recover the COMP and ELEC annotations with perfect accuracy (i.e., *F*_max_ of 1) may be due to inherent errors in these predicted annotations (Škunca *et al.*, 2012). Nevertheless, to further improve our accuracy in recovering these annotations, it may be valuable to integrate phylogenetic information (Jain and Kihara, 2018) or predicted protein secondary structure (Zhang *et al.*, 2018) into the multi-species networks. Finally, while we focused on bacteria, our methods apply generally to any set of related species. Our results point to the feasibility and promise of multi-species, genomewide gene function prediction, especially as more experimental data and annotations become available for diverse organisms.

## 5 Acknowledgements

The research is supported by NSF grants CCF-1617678, DBI-1759858, and MCB-1817736, and by the Office of the Director of National Intelligence (ODNI), Intelligence Advanced Research Projects Activity (IARPA), via the Army Research Office (ARO) under cooperative Agreement Number [W911NF-17-2-0105]. The views and conclusions contained herein are those of the authors and should not be interpreted as necessarily representing the official policies or endorsements, either expressed or implied, of the ODNI, IARPA, ARO, or the U.S. Government. The U.S. Government is authorized to reproduce and distribute reprints for Governmental purposes notwithstanding any copyright annotation thereon.

