## Supplementary file for "Accurate and Efficient Gene Function Prediction using a Multi-Bacterial Network"

### Supplementary File for Accurate and Efficient Network-based Gene Function Prediction using a Multi-Bacterial Network

#### Supplementary Figures

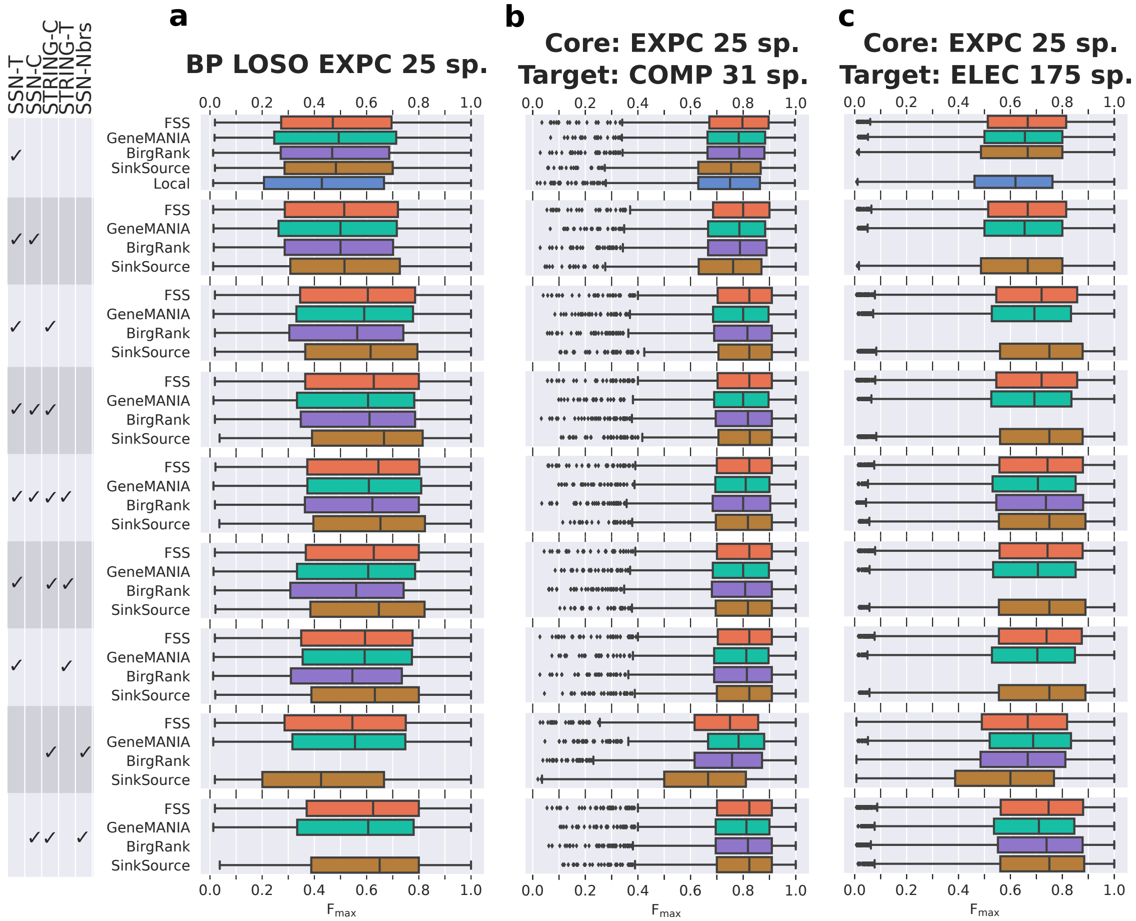


Figure S1: Boxplots of the $F_{\max}$ values for the BP (**a**) EXPC LOSO, (**b**) COMP, (**c**) and ELEC evaluations across nine networks and five algorithms. In every boxplot, the box denotes the 1st and 3rd quartiles, with whiskers extending to the largest value less than 1.5 times the interquartile range.

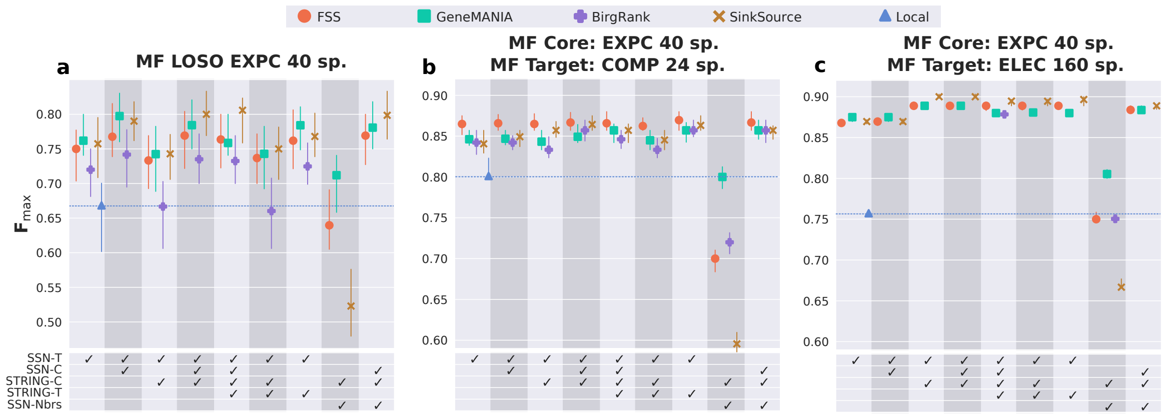


Figure S2: Comparison of $F_{max}$ results for MF EXPC LOSO evaluation, as well as the evaluation of species with only COMP and ELEC annotations across five algorithms and nine networks. (**a-c**) Median $F_{max}$ as well as the 95% confidence interval of the median, estimated using 1,000 bootstrapped samples of the data. Boxplots of the full $F_{max}$ distributions appear in Figure S3. (**a**) Results for EXPC LOSO evaluation; 40 species and 126 MF terms (218 total pairs). (**b**) Results for evaluation of species with COMP and no EXPC; 24 species and 140 MF terms (750 total pairs). (**c**) Results for evaluation of species with ELEC and no EXPC; 160 species and 181 MF terms (11,057 total pairs). We limited algorithms to a maximum of 72 hours of running time, which explains the missing points for BirgRank.

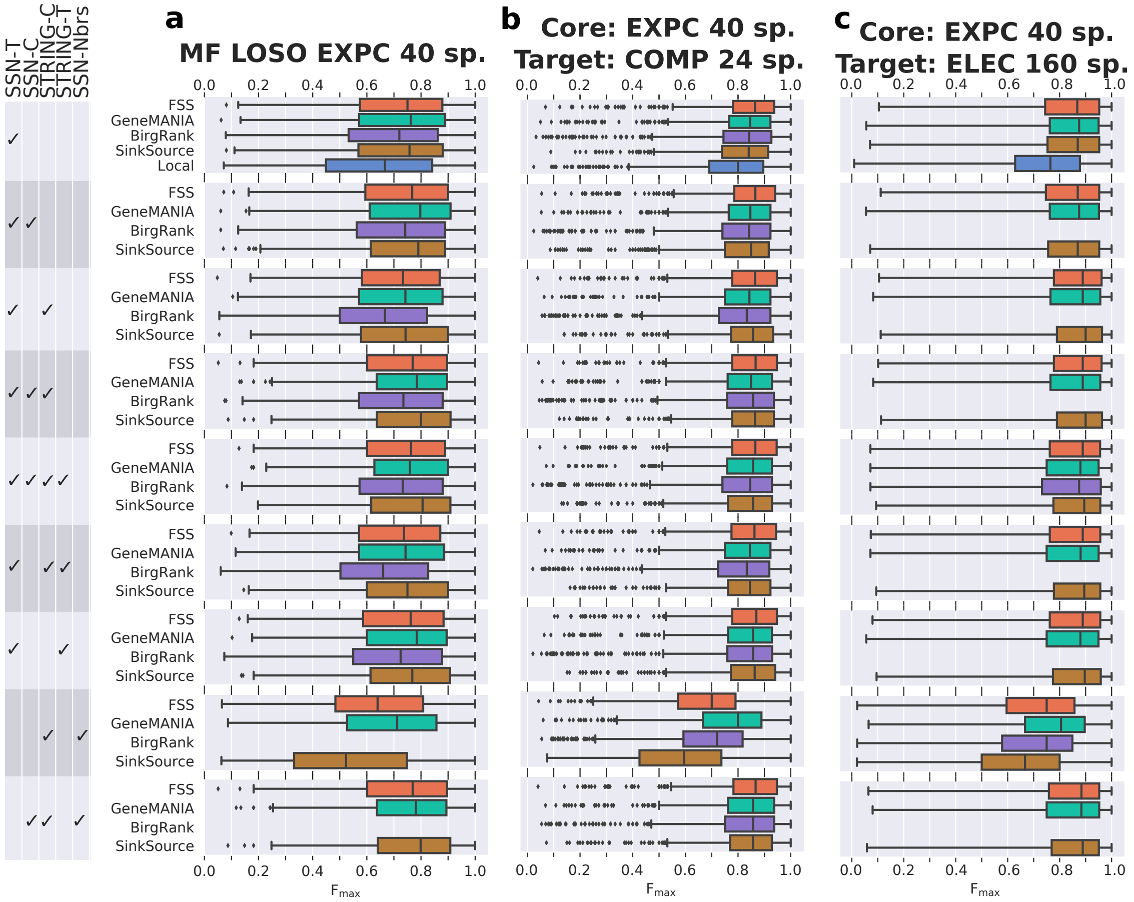


Figure S3: Boxplots of the $F_{max}$ values for the MF (**a**) EXPC LOSO, (**b**) COMP, (**c**) and ELEC evaluations across nine networks and five algorithms. extending to the largest value less than 1.5 times the interquartile range. Note that in (**c**) we extended the whiskers to include all values.

#### Supplementary Tables

| NCBI Taxon ID | Species Name | # BP Ann. | # Genes BP Ann. | # Genes BP EXPC+COMP | # Genes BP EXPC |
|---|---|---|---|---|---|
| 83333 | *E. coli* | 15,479 | 3,397 | 2,511 | 2,348 |
| 83332 | *M. tuberculosis* | 9,005 | 2,363 | 1,659 | 862 |
| 208964 | *P. aeruginosa* | 12,005 | 3,637 | 2,090 | 757 |
| 224308 | *B. subtilis* | 9,115 | 2,620 | 984 | 164 |
| 246196 | *M. smegmatis* | 7,064 | 3,140 | 67 | 56 |
| 99287 | *S. typhimurium* | 10,744 | 3,126 | 1,273 | 52 |
| 190650 | *C. vibrioides* | 4,948 | 1,776 | 53 | 51 |
| 196627 | *C. glutamicum* | 4,061 | 1,504 | 50 | 47 |
| 1111708 | *S. sp. PCC 6803* | 5,910 | 1,764 | 659 | 42 |
| 246200 | *R. pomeroyi* | 7,584 | 3,668 | 512 | 37 |
| 100226 | *S. coelicolor* | 9,611 | 3,793 | 1,236 | 30 |
| 176299 | *A. fabrum* | 6,437 | 2,783 | 61 | 30 |
| 169963 | *L. monocytogenes* | 5,606 | 1,701 | 648 | 25 |
| 160488 | *P. putida* | 7,812 | 2,996 | 28 | 25 |
| 243277 | *V. cholerae* | 11,356 | 3,695 | 1,911 | 24 |
| 93061 | *S. aureus* | 5,465 | 1,562 | 620 | 24 |
| 632 | *Y. pestis* | 7,958 | 2,462 | 548 | 21 |
| 85962 | *H. pylori* | 3,380 | 896 | 389 | 20 |
| 220664 | *P. fluorescens* | 10,028 | 4,848 | 36 | 20 |
| 300852 | *T. thermophilus* | 3,506 | 1,189 | 19 | 19 |
| 266834 | *R. meliloti* | 7,738 | 3,318 | 20 | 18 |
| 242231 | *N. gonorrhoeae* | 3,650 | 1,128 | 18 | 18 |
| 192222 | *C. jejuni* | 3,182 | 997 | 18 | 18 |
| 243230 | *D. radiodurans* | 4,711 | 1,506 | 577 | 15 |
| 272562 | *C. acetobutylicum* | 4,913 | 1,827 | 15 | 12 |

Table S1: Overview of the number of BP annotations in each of the 25 BP core species. The column "# BP Ann." shows the number of such annotations in the GAF file with an EXPC, COMP or ELEC evidence code. The columns with "# Genes" in the title show how many genes have at least one BP annotation with the given evidence codes, where the "# Genes BP Ann." column has the EXPC, COMP and ELEC evidence codes.

| NCBI Taxon ID | Species Name | # MF Ann. | # Genes MF Ann. | # Genes MF EXPC+COMP | # Genes MF EXPC |
|---|---|---|---|---|---|
| 83333 | *E. coli* | 23001 | 3248 | 2655 | 2448 |
| 83332 | *M. tuberculosis* | 12646 | 2463 | 1639 | 678 |
| 208964 | *P. aeruginosa* | 15694 | 3607 | 2139 | 519 |
| 224308 | *B. subtilis* | 12601 | 2628 | 1278 | 268 |
| 882 | *D. vulgaris* | 7381 | 1753 | 225 | 224 |
| 99287 | *S. typhimurium* | 14651 | 3031 | 1474 | 96 |
| 85962 | *H. pylori* | 4913 | 925 | 524 | 96 |
| 1111708 | *S. sp. PCC 6803* | 8472 | 1844 | 882 | 83 |
| 246196 | *M. smegmatis* | 13475 | 3967 | 70 | 57 |
| 246200 | *R. pomeroyi* | 10136 | 2709 | 549 | 56 |
| 243274 | *T. maritima* | 5897 | 1186 | 581 | 46 |
| 300852 | *T. thermophilus* | 5781 | 1354 | 46 | 43 |
| 243277 | *V. cholerae* | 12314 | 2353 | 1980 | 42 |
| 100226 | *S. coelicolor* | 16196 | 4634 | 1680 | 40 |
| 176299 | *A. fabrum* | 10483 | 2980 | 52 | 39 |
| 160488 | *P. putida* | 12351 | 3208 | 43 | 38 |
| 192222 | *C. jejuni* | 4857 | 1052 | 36 | 35 |
| 83334 | *E. coli* | 12970 | 3048 | 64 | 31 |
| 632 | *Y. pestis* | 11075 | 2369 | 629 | 26 |
| 196627 | *C. glutamicum* | 7016 | 1736 | 30 | 25 |
| 211586 | *S. oneidensis* | 12243 | 2541 | 1460 | 23 |
| 266834 | *R. meliloti* | 13420 | 3594 | 27 | 22 |
| 223283 | *P. syringae* | 11222 | 3004 | 369 | 21 |
| 190650 | *C. vibrioides* | 7531 | 2051 | 25 | 21 |
| 272623 | *L. lactis* | 5703 | 1343 | 28 | 21 |
| 93061 | *S. aureus* | 7719 | 1609 | 744 | 20 |
| 272562 | *C. acetobutylicum* | 8236 | 2215 | 25 | 19 |
| 224324 | *A. aeolicus* | 5525 | 1061 | 539 | 19 |
| 623 | *S. flexneri* | 11057 | 2679 | 43 | 18 |
| 226186 | *B. thetaiotaomicron* | 10087 | 2773 | 1177 | 18 |
| 190485 | *X. campestris* | 10237 | 2444 | 1091 | 17 |
| 103690 | *N. sp. PCC 7120* | 9112 | 2523 | 25 | 15 |
| 71421 | *H. influenzae* | 7222 | 1278 | 663 | 15 |
| 1140 | *S. elongatus* | 6269 | 1443 | 15 | 14 |
| 272943 | *R. sphaeroides* | 10160 | 2488 | 18 | 13 |
| 243276 | *T. pallidum* | 2680 | 548 | 15 | 13 |
| 196600 | *V. vulnificus* | 10323 | 2562 | 16 | 12 |
| 272942 | *R. capsulatus* | 9337 | 2156 | 16 | 12 |
| 220668 | *L. plantarum* | 7781 | 1956 | 13 | 11 |
| 122586 | *N. meningitidis* | 6334 | 1187 | 606 | 11 |

Table S2: Overview of the number of MF annotations in each of the 40 MF core species. The column "# MF Ann." shows the number of such annotations in the GAF file with an EXPC, COMP or ELEC evidence code. The columns with "# Genes" in the title show how many genes have at least one MF annotation with the given evidence codes, where the "# Genes MF Ann." column has the EXPC, COMP and ELEC evidence codes.

| SSN E-value cutoff | STRING cutoff | # Nodes | # Edges |
|---|---|---|---|
| $1 \times 10^{-25}$ | - | 74,368 | 590,272 |
| $1 \times 10^{-15}$ | - | 80,354 | 756,802 |
| $1 \times 10^{-6}$ | - | 86,257 | 1,000,169 |
| 0.1 | - | 89,872 | 1,219,556 |
| 5 | - | 92,787 | 1,317,828 |
| 20 | - | 95,472 | 1,369,983 |
| 0.1 | 900 | 91,785 | 1,395,330 |
| 0.1 | 700 | 96,107 | 1,596,336 |
| 0.1 | 400 | 98,849 | 2,228,049 |
| 0.1 | 150 | 98,891 | 7,502,222 |

Table S3: Network sizes for various BLAST E-value and STRING cutoffs for the 25 core species on the SSN-T+SSN-C network (first six rows) and the full network (last four rows; SSN-T+SSN-C+STRING-C+STRING-T).

| | Target: COMP 31 sp. | | Target: ELEC 175 sp. | |
|---|---|---|---|---|
| Network Combination | # Nodes | # Edges | # Nodes | # Edges |
| SSN-T | 170,097 | 2,217,014 | 674,888 | 15,459,229 |
| SSN-T+SSN-C | 177,439 | 3,436,570 | 676,084 | 16,678,785 |
| SSN-T+STRING-C | 179,335 | 2,602,036 | 677,310 | 15,844,251 |
| SSN-T+SSN-C+STRING-C | 182,029 | 3,813,350 | 678,088 | 17,055,565 |
| SSN-T+SSN-C+STRING-C+STRING-T | 188,347 | 4,077,538 | 716,367 | 19,028,238 |
| SSN-T+STRING-C+STRING-T | 185,653 | 2,866,224 | 715,589 | 17,816,924 |
| SSN-T+STRING-T | 176,415 | 2,481,202 | 713,167 | 17,431,902 |
| STRING-C,SSN-Nbrs | 76,814 | 385,022 | 76,814 | 385,022 |
| SSN-C+STRING-C,SSN-Nbrs | 96,107 | 1,596,336 | 96,107 | 1,596,336 |

Table S4: Network sizes for the nine network combinations when connecting the 25 BP core species to 31 and 175 species for the COMP and ELEC evaluations, respectively. Note that for the combinations with SSN-Nbrs, we show only the size of the core network

| | A | B | C | D | E | F | G | H | I |
|---|---|---|---|---|---|---|---|---|---|
| A | | 0.3 | $\mathbf{8.5 \times 10^{-8}}$ | $\mathbf{3.5 \times 10^{-10}}$ | $\mathbf{1.2 \times 10^{-10}}$ | $\mathbf{1.2 \times 10^{-8}}$ | $\mathbf{2.0 \times 10^{-7}}$ | **0.02** | $\mathbf{9.5 \times 10^{-11}}$ |
| B | 1.0 | | $\mathbf{4.1 \times 10^{-5}}$ | $\mathbf{1.2 \times 10^{-6}}$ | $\mathbf{5.3 \times 10^{-8}}$ | $\mathbf{1.1 \times 10^{-6}}$ | $\mathbf{3.6 \times 10^{-5}}$ | 0.14 | $\mathbf{8.6 \times 10^{-8}}$ |
| C | 1.0 | 1.0 | | 0.28 | **0.01** | 0.28 | 0.71 | 1.0 | 0.22 |
| D | 1.0 | 1.0 | 1.0 | | 0.37 | 0.80 | 1.0 | 1.0 | 0.88 |
| E | 1.0 | 1.0 | 1.0 | 1.0 | | 1.0 | 1.0 | 1.0 | 1.0 |
| F | 1.0 | 1.0 | 1.0 | 1.0 | 0.18 | | 1.0 | 1.0 | 0.75 |
| G | 1.0 | 1.0 | 1.0 | 0.15 | **0.01** | 0.18 | | 1.0 | 0.08 |
| H | 1.0 | 1.0 | $\mathbf{1.2 \times 10^{-3}}$ | $\mathbf{1.4 \times 10^{-5}}$ | $\mathbf{2.1 \times 10^{-7}}$ | $\mathbf{2.1 \times 10^{-5}}$ | $\mathbf{1.4 \times 10^{-3}}$ | | $\mathbf{3.3 \times 10^{-5}}$ |
| I | 1.0 | 1.0 | 1.0 | 1.0 | 0.42 | 1.0 | 1.0 | 1.0 | |

Table S5: Benjamini-Hochberg-corrected $q$-values for the one-sided Wilcoxon signed-rank test (improvement of column over row) corresponding to the FastSinkSource results in Figure 3(a). Bold text highlights $q$-values $\leq$ 0.05. Letters represent the network combinations as follows: A: SSN-T; B: SSN-T+SSN-C; C: SSN-T+STRING-C; D: SSN-T+SSN-C+STRING-C; E: SSN-T+SSN-C+STRING-C+STRING-T; F: SSN-T+STRING-C+STRING-T; G: SSN-T+STRING-T; H: STRING-C,SSN-Nbrs; I: SSN-C+STRING-C,SSN-Nbrs;

### Supplementary Text

#### S1 Properties of the FastSinkSource Algorithm

To keep this section self-contained, we start by reformulating the FastSinkSource algorithm. Recall that the input to FastSinkSource is a weighted undirected graph $G = (V, E, w)$, where $w_{uv} \geq 0$ represents the weight of the edge $(u, v)$. In addition, for a GO term $\tau$, we have a partition of the nodes in $V$ into three sets: $V^+$, $V^-$ and $V^0$ positive, negative, and unknown examples, respectively. We also have a parameter $0 < \alpha < 1$ in the input. Note that we do not permit $\alpha = 1$ here since some of our lemmas and proofs do not apply to this value.

We fix the score of every positive example at 1 and for every negative example at 0. The goal is to compute a score $s(u)$ between 0 and 1 for each node $u$ that is an unknown example by propagating the positive and negative labels across $G$ using the following equation:

$$s(u) = \frac{\alpha \sum\limits_{v \in N(u)} w_{uv} s(v)}{d(u)} = \frac{\alpha \sum\limits_{v \in N^0(u)} w_{uv} s(v)}{d(u)} + f(u) \tag{1}$$

where $N(u)$ is the set of neighbors of node $u$, $N^0(u)$ is the subset of neighbors of node $u$ that are unknown examples, $d(u)$ is the weighted degree of node $u$ in $G$, and

$$f(u) = \frac{\alpha \sum\limits_{v \in N^+(u)} w_{uv}}{d(u)},$$

where $N^+(u)$ is the set of neighbors of $u$ that are positive examples. Note that $f(u) \geq 0$ is a constant for every node $u$.

We define $\mathbf{s}$ to be a $|V^0| \times 1$ vector formed by the node scores of all unknown examples and $\mathbf{f}$ to be a $|V^0| \times 1$ constant vector in which the $u$th element is $f(u)$. Further, we let the $|V^0| \times |V^0|$ matrix $\mathbf{P}$ contain the degree-normalized edge weights in $G$ among pairs of nodes in $V^0$, i.e., $P_{uv} = w_{uv}/d(u)$ for every pair of unknown examples $(u, v)$. Then for every unknown example $u$, we can combine the score equations (1) into a single linear system:

$$\mathbf{s} = \alpha \mathbf{P} \mathbf{s} + \mathbf{f}. \tag{2}$$

We use power iterations to solve this system of linear equations. Let $s(u, i)$ denote the score computed for node $u$ after $i$ iterations and let $\mathbf{s}^{(i)}$ denote the vector formed by these scores after $i$ iterations. We initialize by $s(u, 0) = 0$ for every node $u$ in $G$, i.e., we set $\mathbf{s}^{(i)} = 0$. For every node $u$, we compute its score in iteration $i + 1$ using the equation

$$s(u, i + 1) = \alpha \sum_{v \in N^0(u)} P_{uv} s(v, i) + f(u) \tag{3}$$

In other words, we have that

$$\mathbf{s}^{(i+1)} = \alpha \mathbf{P} \mathbf{s}^{(i)} + \mathbf{f}. \tag{4}$$

We first prove that the score for every node $u$ does not decrease from one iteration to the next, is a lower bound on $s(u)$, and reaches this value in the limit. We say that a vector is *non-negative* if every element in it is non-negative. Given two vectors $\mathbf{x}$ and $\mathbf{y}$, we use $\mathbf{x} \leq \mathbf{y}$ to denote that $\mathbf{y} - \mathbf{x}$ is non-negative.

**Lemma S1.1.** *For every node $u \in V$, $s(u, i) \leq s(u, i + 1) \leq s(u)$ for every $i \geq 1$.*

*Proof.* Combining Equation (3) with the fact that no entry in $\mathbf{P}$ is negative, we see that

$$s(u, i + 1) - s(u, i) = \alpha \sum_{v \in N^0(u)} P_{uv} \big( s(v, i) - s(v, i - 1) \big) \geq 0,$$

Moreover, by applying induction to eq. (4), we can prove that

$$\mathbf{s}^{(i)} = \sum_{j=0}^{i} (\alpha \mathbf{P})^j \mathbf{f} \ \leq \ \sum_{j=0}^{\infty} (\alpha \mathbf{P})^j \mathbf{f}, \tag{5}$$

where the last inequality follows from the fact that every entry in any power of $\mathbf{P}$ is non-negative. We can prove that for every $j \geq 1$, every row sum in $\mathbf{P}^j$ is bounded from above by one. Therefore, in combination with $0 < \alpha < 1$, we have $\lim_{j\to\infty}(\alpha\mathbf{P})^j = 0$. We can further show that $\mathbf{I} - \alpha\mathbf{P}$ is invertible and that this inverse equals the infinite sum on the right-hand size of Equation (5). Moreover, since $\mathbf{I} - \alpha\mathbf{P}$ is invertible, we can solve eq. (2) for $\mathbf{s}$ to obtain $\mathbf{s} = (\mathbf{I} - \alpha\mathbf{P})^{-1}\mathbf{f}$. We conclude that for every $i \geq 1, \mathbf{s}^{(i)} \leq \mathbf{s}$, which proves the inequality on the right-hand side of the lemma. □

This proof implies the following corollary to the lemma.

**Corollary S1.1.** *For every node $u \in V$, $\lim_{i\to\infty} s(u,i) = s(u)$.*

We now prove an upper bound on $s(u,i)$ for every node $u$. In the following lemma and in its proof, we use $\|\mathbf{f}\|$ to denote the largest value in $\mathbf{f}$. For a matrix $\mathbf{A}$, we use $\|\mathbf{A}\|$ to denote the largest row sum in $A$. Note that $\|\mathbf{P}\| = 1$ and that $\|\mathbf{P}\mathbf{s}\| \leq \|\mathbf{P}\|\|\mathbf{s}\|$.

**Lemma S1.2.** *For every node $u \in V$ and for every $i \geq 1$,*

$$s(u, i+1) \leq s(u,i) + \alpha^i\|\mathbf{f}\| \text{ and}$$
$$s(u) \leq s(u,i) + \frac{\alpha^i\|\mathbf{f}\|}{(1-\alpha)}$$

*Proof.* For every $i \geq 1$, Equation (4) implies that

$$\mathbf{s}^{(i+1)} - \mathbf{s}^{(i)} = \alpha\mathbf{P}(\mathbf{s}^{(i)} - \mathbf{s}^{(i-1)})$$

By computing the maximum value on both sides, we have

$$\begin{aligned}\|\mathbf{s}^{(i+1)} - \mathbf{s}^{(i)}\| &= \|\alpha\mathbf{P}(\mathbf{s}^{(i)} - \mathbf{s}^{(i-1)})\| \\ &\leq \|\alpha\mathbf{P}\| \cdot \|\mathbf{s}^{(i)} - \mathbf{s}^{(i-1)}\| \\ &= \alpha\|\mathbf{s}^{(i)} - \mathbf{s}^{(i-1)}\|\end{aligned}$$

By combining induction with this inequality, we can prove that

$$\|\mathbf{s}^{(i+1)} - \mathbf{s}^{(i)}\| \leq \alpha^i\|\mathbf{s}^{(1)} - \mathbf{s}^{(0)}\| = \alpha^i\|\mathbf{f}\| \tag{6}$$

Thus, we have shown that the for every node, the difference between its scores in iterations $i+1$ and $i$ decreases geometrically with $i$, thereby proving the first inequality in the lemma. To complete the proof by demonstrating the second inequality, we need to relate the node's score at convergence to its score after $i$ iterations. We do so by proving an upper bound on how much the node's score can increase after $i$ iterations.

For every integer $m > i$, we have

$$\begin{aligned}\|\mathbf{s}^{(m)} - \mathbf{s}^{(i)}\| &= \|\sum_{j=i}^{m-1}(\mathbf{s}^{(j+1)} - \mathbf{s}^{(j)})\| \\ &\leq \sum_{j=i}^{m-1}\|\mathbf{s}^{(j+1)} - \mathbf{s}^{(j)}\| \\ &\leq \sum_{j=i}^{m-1}\alpha^j\|\mathbf{f}\| \qquad \text{by Equation (6)} \\ &= \alpha^i\|\mathbf{f}\|\sum_{j=0}^{m-i-1}\alpha^j = \alpha^i\|\mathbf{f}\|\left(\frac{1-\alpha^{m-i}}{1-\alpha}\right)\end{aligned}$$

Since $\lim_{m\to\infty} s(u,m) = s(u)$ by Corollary S1.1, we have that

$$\begin{aligned}\|\mathbf{s} - \mathbf{s}^{(i)}\| &= \lim_{m\to\infty}\|\mathbf{s}^{(m)} - \mathbf{s}^{(i)}\| \\ &\leq \alpha^i\|\mathbf{f}\|\lim_{m\to\infty}\left(\frac{1-\alpha^{m-i}}{1-\alpha}\right) \leq \frac{\alpha^i\|\mathbf{f}\|}{(1-\alpha)},\end{aligned}$$

which completes the proof. □

We now consider how the node scores after each iteration change when we decrease the value of $\alpha$. We first prove that as $\alpha$ decreases, the score for each node after $i$ iterations also decreases. Then we prove that as $\alpha$ decreases, the difference in node scores from one iteration to the next also decreases, which means that fewer iterations are necessary for convergence. We introduce an additional piece of notation here. Let $s_\beta(u,i)$ be the score of node $u$ after $i$ iterations of SinkSource for a given GO term and with the value of $\alpha$ set to $\beta$.

**Lemma S1.3.** *Let $0 < \beta < \gamma \le 1$. Then for every node $u \in V$ and for every integer $i \ge 1$, $s_\beta(u,i) \le s_\gamma(u,i)$.*

*Proof.* We prove the lemma by induction. To prove the base that $s_\beta(u,1) \le s_\gamma(u,1)$, we observe that

$$\begin{aligned} s_\beta(u,1) &= \beta \sum_{v \in N^0(u)} P_{uv} s_\beta(v,0) + \beta \sum_{v \in N^+(u)} p_{uv}, \text{ by Equation (3)} \\ &= \beta \sum_{v \in N^+(u)} p_{uv}, \text{ since } s_\beta(v,0) = \text{ for every node } v \in V \end{aligned}$$

Similarly,

$$s_\gamma(u,1) = \gamma \sum_{v \in N^+(u)} p_{uv}$$

Since $\beta < \gamma$, we have

$$s_\beta(u,1) \le s_\gamma(u,1)$$

The inductive hypothesis is that for some integer $i - 1 \ge 1$, $s_\beta(u, i-1) \le s_\gamma(u, i-1)$ for every node $u \in V$. We have that

$$\begin{aligned} s_\beta(u,i) &= \beta \sum_{v \in N^0(u)} P_{uv} s_\beta(v, i-1) + \beta \sum_{v \in N^+(u)} p_{uv} \\ &\le \beta \sum_{v \in N^0(u)} p_{uv} s_\beta(v, i-1) + \gamma \sum_{v \in N^+(u)} p_{uv}, \text{ since } s_\beta(u,1) \le s_\gamma(u,1) \\ &\le \beta \sum_{v \in N^0(u)} p_{uv} s_\gamma(v, i-1) + \gamma \sum_{v \in N^+(u)} p_{uv}, \text{ by the inductive hypothesis} \\ &< \gamma \sum_{v \in N^0(u)} p_{uv} s_\gamma(v, i-1) + \gamma \sum_{v \in N^+(u)} p_{uv}, \text{ since } \beta < \gamma \\ &= s_\gamma(u,i) \end{aligned}$$

thus proving the statement for $i$ and completing the proof. □

We can also show that the difference between a node's score in iterations $i$ and $i - 1$ itself decreases as the value of $\alpha$ becomes smaller.

**Lemma S1.4.** *Let $0 < \beta < \gamma \le 1$. Then for every node $u \in V$ and for every integer $i \ge 1$,*

$$s_\beta(u,i) - s_\beta(u,i-1) \le s_\gamma(u,i) - s_\gamma(u,i-1).$$

*Proof.* Consider any integer $i \ge 1$. By Equation (3), we have

$$\begin{aligned} s_\beta(u,i) - s_\beta(u,i-1) &= \beta \sum_{v \in N^0(u)} P_{uv} s_\beta(v,i-1) - \beta \sum_{v \in N^0(u)} P_{uv} s_\beta(v,i-2), \\ &\quad \text{since the contributions from the positively-labeled neighbors cancel each other} \\ &< \gamma \sum_{v \in N^0(u)} P_{uv} s_\beta(v,i-1) - \gamma \sum_{v \in N^0(u)} P_{uv} s_\beta(v,i-2), \text{ since } \beta < \gamma \\ &\le \gamma \sum_{v \in N^0(u)} P_{uv} s_\gamma(v,i-1) - \gamma \sum_{v \in N^0(u)} P_{uv} s_\gamma(v,i-2), \text{ by Lemma S1.3} \\ &= s_\gamma(u,i) - s_\gamma(u,i-1), \end{aligned}$$

thus completing the proof. □

#### S2 Other Algorithms

**Local.** Each gene's score for a GO term is the weighted average of the scores of its neighbors, where the positive examples for a term get a score of 1. Local mimics a basic procedure for assigning GO term annotations to a query gene $q$: use BLAST to determine if $q$'s sequence is similar to that of a gene $q'$ in another organism and then transfer the annotations of $q'$ to $q$.

**GeneMANIA (Mostafavi *et al.*, 2008).** Given a weighted, undirected network $G = (V, E, w)$, and a label vector $\mathbf{y}$ where $y(u)$ represents the prior evidence for gene $u$ having a function of interest, this algorithm computes a discriminant score $s(u)$ between $-1$ and 1 to each gene $u$ in the network, which we can threshold to classify the genes. The value of $y(u)$ is 1 or -1 if $u$ is a positive or negative example, respectively. Let the number of positive examples be $n^+$ and the number of negative example be $n^-$. If $u$ is an unknown example, then $y(u) = \frac{n^+ - n^-}{n^+ + n^-}$, the mean of the labels of the labeled nodes. To compute the vector containing the discriminant scores for each node, we solve the following optimization problem:

$$\mathbf{s} = \arg\min_{\mathbf{s}} \left( \sum_{u \in V} (s(u) - y(u))^2 + \sum_{(u,v) \in E} w_{uv} (s(u) - s(v))^2 \right), \tag{7}$$

where the minimization ranges over all vectors in $\mathbb{R}^m$ and $m$ is the number of nodes in the graph. Let $\mathbf{W} \in \mathbb{R}^{m \times m}$ denote the adjacency matrix of $G$, and $\mathbf{P} = \mathbf{D}^{-1/2}\mathbf{W}\mathbf{D}^{-1/2}$ denote the normalized network, where $\mathbf{D}$ is a diagonal matrix with $\mathbf{D}_{uu} = \sum_v w_{uv}$. To minimize the sum in Equation (7), we solve the linear system of equations $(\mathbf{I} + \mathbf{D} - \mathbf{P})\mathbf{s} = \mathbf{y}$.

**BirgRank (Jiang *et al.*, 2017).** While BirgRank is also a network propagation algorithm, it has significant differences from SinkSource and GeneMANIA. As we describe below, the first difference is that it directly incorporates the GO hierarchy into the diffusion process. Second, BirgRank propagates labels on a gene-by-gene basis, meaning it makes all function predictions for a single gene simultaneously. Third, it does not incorporate negative examples.

BirgRank utilizes three datasets: i) a weighted, undirected network $G$ represented as the column normalized adjacency matrix $\mathbf{W} \in \mathbb{R}^{m \times m}$ where $m$ is the number of genes; ii) a set of gene-function annotations represented in a binary matrix $\mathbf{R} \in \mathbb{R}^{m \times n}$ where $n$ is the number of functions, and $R_{ij} = 1$ if gene $i$ is annotated by function $j$, and is 0 otherwise; and iii) a function hierarchy represented by a binary matrix $\mathbf{H} \in \mathbb{R}^{n \times n}$ where $H_{ij} = 1$ if the term $i$ is a child of term $j$, and is 0 otherwise. BirgRank combines these three matrices into a single matrix in order to incorporate the hierarchy $\mathbf{H}$ into the propagation. BirgRank performs random walks with restarts (RWR) controlled by four parameters: (a) $\alpha$ is the standard parameter for the restart probability, (b) $\mu$ controls the proportion of transition along edges in $\mathbf{W}$ versus along edges in $\mathbf{R}$ (that connect genes in $G$ to the terms annotating them in $H$), (c) $\theta$ controls the probability of restart from a given gene versus the probability of restart from the functions annotated to it, and (d) $\lambda$ controls the direction of transitions within the hierarchy (i.e., up or down) with $\mathbf{H}^* = \lambda\mathbf{H} + (1 - \lambda)\mathbf{H}^T$.

To compute the prediction scores, BirgRank solves the following system of linear equations:

$$\left( \begin{bmatrix} \mathbf{I}_m & 0 \\ 0 & \mathbf{I}_n \end{bmatrix} - \alpha \overline{\begin{bmatrix} \mu\mathbf{W} & 0 \\ (1-\mu)\mathbf{R}^T & \mathbf{H}^* \end{bmatrix}} \right) \begin{bmatrix} \mathbf{X}_W \\ \mathbf{X}_H \end{bmatrix} = (1-\alpha) \begin{bmatrix} \theta\mathbf{I}_m \\ (1-\theta)\mathbf{R}^T \end{bmatrix}, \tag{8}$$

where the bar over the block matrix indicates the the whole matrix is column normalized, and $I_n$ and $I_m$ are $n \times n$ and $m \times m$ identity matrices, respectively. Each column vector $\mathbf{v}$ in $\mathbf{X}_W \in \mathbb{R}^{m \times m}$ contains the RWR scores for a single gene, i.e., $\mathbf{X}_{W uv}$ is the probability that a random walker who restarts to gene $v$ will visit gene $u$. Each column vector in the lower block of the solution, $\mathbf{X}_H \in \mathbb{R}^{n \times m}$, contains the RWR scores from a single gene to the nodes corresponding to GO terms, i.e., $\mathbf{X}_{H uv}$ is the probability of the random walker who restarts to gene $v$ or to $v$'s annotations in $\mathbf{R}$ will visit the term $u$. We derive the function predictions scores for each gene from $\mathbf{X}_H$.

In the original BirgRank paper (Jiang *et al.*, 2017), the annotation matrix $\mathbf{R}$ contained only the direct annotations. Since we evaluated on a term-by-term basis, we opted to include all propagated annotations in $\mathbf{R}$, i.e., if a gene was annotated to a GO term, we include annotations of that gene to all ancestors of that term in the GO hierarchy.

The original MATLAB implementation of BirgRank solved for $\mathbf{X}_W$ and $\mathbf{X}_H$ directly, i.e., by computing the inverse of the matrix on the left of Equation (8) and multiplying by the vector on the right, thus making predictions for all nodes. We were able to greatly reduce the running time of BirgRank by computing scores for only the left-out positive and negative examples in LOSO validations. To select the parameters for BirgRank, we systematically varied them and performed LOSO evaluation (BP EXPC annotations, SSN+STRING network). We selected the values that gave the highest median $F_{\max}$: $\alpha = 0.95$, $\mu = 0.5$, $\theta = 0.5$, and $\lambda = 0.01$ (Section S4.4).

**Implementation.** We implemented each of these algorithms (SinkSource, FastSinkSource, Local, GeneMANIA, and BirgRank) in Python using the SciPy (v1.1.0) and NumPy (v1.15) libraries. For GeneMANIA and BirgRank, we translated their MATLAB implementations on a line-by-line basis to SciPy sparse matrix operations. We ensured the correctness of our implementations by comparing our output values to those output by the corresponding MATLAB implementation, using the 19-species SSN and BP EXPC annotations. They matched exactly. To solve the GeneMANIA linear system, we utilized the conjugate gradient (CG) solver implemented in the SciPy (v1.1.0) Python package and used SciPy's default tolerance cutoff of $1 \times 10^{-5}$. To compute the scores for BirgRank, we used power iteration and stopped when the maximum difference of node scores between iterations $i$ and $i-1$ (i.e., $\epsilon$) was $\leq 1 \times 10^{-4}$).

### S3 Datasets

#### S3.1 Positive, Negative, and Unknown Examples

We say a given gene $g$ is *directly* annotated to a GO term $t$ if this annotation appears in the annotations (GAF) file. For a given $t$, we defined $g$ as

(a) a *positive example* if $g$ was directly annotated to $t$ or to a descendant of $t$ in the GO DAG,

(b) a *negative example* if $g$ was not directly annotated to $t$ or to an ancestor or descendant of $t$ in the GO DAG, did not share a direct annotation with any of the genes annotated to $t$, and had at least one other annotation, and

(c) an *unknown example* otherwise.

We adopted this definition of negative examples from Youngs *et al.* (Youngs *et al.*, 2013). We used only the "is_a" edge type of the Gene Ontology when defining these positive and negative examples. In several analyses, we restricted our attention to a specific set of evidence codes, e.g., EXPC. In such a situation, we discarded all direct annotations with other evidence codes before computing the different sets of examples.

We utilized the negative examples in different aspects of this work: (a) for algorithms that use negatives examples (SinkSource and GeneMANIA) (Sections S2 and 2.1), (b) for integrating multiple networks, (Section S3.2), and (c) for defining false positives when calculating the $F_{\max}$ during evaluation (Section 2.5).

#### S3.2 Network Integration

We compared two kernel-based network integration methods.

**GMW:** Mostafavi and Morris integrated multiple networks on a GO term-by-GO term basis using ridge regression (Mostafavi *et al.*, 2008). Given $m$ networks represented as adjacency matrices $W_1, ..., W_m$, they combined them using a weighted sum $\mathbf{W}^* = \sum_{i=0}^{m} \mu_i W_i$, where they computed the network weights

$\mu^* = [\mu_1, \mu_2, ..., \mu_m]$ independently for each prediction task (i.e., one set of weights for each GO term) by solving the following constrained linear regression problem:

$$\mu^* = \min_{\mu} (\mathbf{t} - \mathbf{\Omega}\mu)^T (\mathbf{t} - \mathbf{\Omega}\mu),$$
$$\text{s.t. } \mu_i \geq 0 \quad i = \{1, ..., m\}.$$

We define the terms in this equation next. If $n^+$ and $n^-$ are the number of positive and negative examples, respectively, and $n = n^+ + n^-$ is the total number of labelled examples, then the target vector $\mathbf{t}$ contains $(n^+)^2 + n^+n^-$ entries, one for each positive-positive example pair and one for each positive-negative example pair. The value in $\mathbf{t}$ is $\frac{n^-}{n}^2$ for each positive-positive example pair and $-\frac{n^+n^-}{n^2}$ for each positive-negative example pairs. The matrix $\Omega$ has $m$ columns, one for each of the networks and $(n^+)^2 + n^+n^-$ rows, corresponding to the positive-positive and positive-negative pairs. The resulting vector $\mu^*$ contains the relative importance of each network, based on how well its edge weights match the positive pairs versus the positive-negative pairs.

**SWSN:** Mostafavi and Morris extended their method to compute optimal weights for multiple related GO terms simultaneously (Simultaneous Weights, SW) (Mostafavi and Morris, 2010). A drawback of this approach was that it treated all non-positive examples as negative examples. Youngs *et al.* later modified the SW method to be able to use specific negative examples for each GO term (Simultaneous Weights with Specific Negatives, SWSN) (Youngs *et al.*, 2013). The SWSN method assigns network weights by solving the following problem:

$$\mu^* = \min_{\mu} \sum_{c=1}^{h} (\mathbf{t}_c - \mathbf{\Omega}\mu)^T (\mathbf{t}_c - \mathbf{\Omega}\mu),$$

where the index $c$ runs over a set of related GO terms. They found that computing the weights to all GO terms in a particular hierarchy (e.g., BP or MF) performed better than any other grouping of functions. Therefore, we also grouped GO terms by hierarchy to compute the optimal weights for our networks.

### S4 Parameter Selection

#### S4.1 Testing the Effect of BLAST E-value Cutoffs

Here we studied the effect of using different E-value cutoffs to construct the sequence-similarity network (SSN). We tested E-value cutoffs of $1 \times 10^{-25}$, $1 \times 10^{-15}$, $1 \times 10^{-6}$, 0.1, 5, and 20. We repeated the BP and MF EXPC LOSO evaluations for each cutoff. We chose to test the methods SinkSource, GeneMANIA, and Local since they did not have any parameters to vary. We observed a general trend that raising the E-value cutoff resulted in better $F_{\max}$ values. For BP terms, the increase in median $F_{\max}$ leveled off around $1 \times 10^{-6}$, whereas a cutoff of 0.1 resulted in the best or near-best median $F_{\max}$ values for MF (Figure S4a,d). In further analyses, we used an E-value cutoff of 0.1 for the SSN.

#### S4.2 Network Integration

Techniques that utilize multiple complementary data sources have been shown to be more accurate than those that use a single data source (Jiang *et al.*, 2016; Cozzetto *et al.*, 2013; Gligorijević *et al.*, 2018). Therefore, we integrated the SSN with networks from the STRING database (Szklarczyk *et al.*, 2019). However, as the ranges of edge weights in the SSN and the STRING networks differ (1–180 and 150–1,000, respectively), we needed to carefully balance and combine these disparate networks to achieve optimal prediction performance. To accomplish this integration, we tested two methods: term-by-term weighting described in the original GeneMANIA publication (GMW; we use this abbreviation to differentiate the weighting method from the network propagation algorithm) (Mostafavi *et al.*, 2008) and Simultaneous Weights with Specific Negatives (SWSN) (Youngs *et al.*, 2013) which weights networks using multiple related terms simultaneously. Each

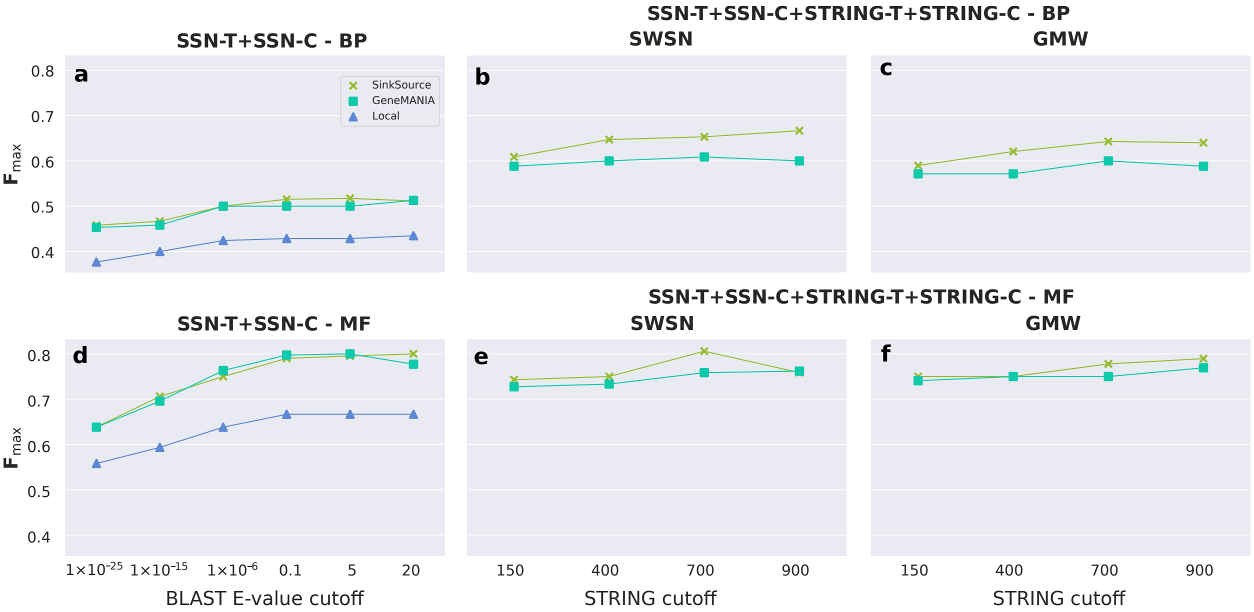


Figure S4: Comparison of the median $F_{\max}$ from the LOSO evaluation of EXPC annotations using various BLAST E-value cutoffs (**a**, **d**) and STRING cutoffs (**b**, **c**, **e**, **f**). Related to Figure 3. (**b**, **e**) SWSN and (**c** and **f**) GMW integration of the STRING networks with the SSN (E-value $< 0.1$). Here we used the full SSN and STRING networks (i.e., SSN-T+SSN-C and SSN-T+SSN-C+STRING-T+STRING-C).

of these methods computes an optimal weight for each network in order to maximize the total weight of the edges between pairs of positively labeled genes while minimizing the total weight of the edges between genes of opposite labels. For the SWSN method, we used all terms of a given hierarchy as this was shown to perform the best in a previous evaluation (Youngs *et al.*, 2013). Note that since BirgRank makes predictions for all GO terms for a given gene simultaneously, we could use only SWSN for it. In addition to deciding between GMW and SWSN, we also considered how to select a cutoff on the edge weights in the STRING networks. To this end, we used low, medium, high, and very high stringency cutoffs (150, 400, 700, and 900, respectively) on the edge weights of the STRING networks and integrated each of these networks with the SSN. We evaluated these networks using BP and MF EXPC LOSO validation on the full network (SSN-T+SSN-C+STRING-T+STRING-C), and tested only SinkSource and GeneMANIA, since they did not have parameters to vary, and Local can use only the SSN-T edges. Note that for each species we left out, we weighted the networks using only annotations of the other organisms we retained (i.e., 24 and 39 for BP and MF, respectively).

For BP and MF GO terms, we found that both algorithms achieved the highest median $F_{\max}$ with either the high or very high stringency cutoffs (Figure S4c). While the performance on the networks generated by GMW and SWSN were comparable, SWSN was faster than GMW (6.4 minutes vs 14.4 minutes for SWSN and GMW, respectively). An additional benefit of SWSN is that the resulting edge weights do not change from one GO term to another. Therefore, we chose to use SWSN with a stringency cutoff of 700 to integrate the networks.

#### S4.3 Assessing the Trade-off between Accuracy and Speed for FastSinkSource

FastSinkSource converges faster, i.e., with a smaller number of iterations, as we decrease the parameter $\alpha$ (see proof in Section S1). Additionally, for a fixed $\alpha$, FastSinkSource terminates faster as we decrease the number of iterations. Therefore, we sought to test the trade-off between accuracy and speed by varying either $\alpha$ or the number of iterations before we stopped FastSinkSource. For this analysis, we performed LOSO validation using the full network (i.e., SSN-T+SSN-C+STRING-T+STRING-C).

We ran FastSinkSource for eight different values of $\alpha$, namely, 1, 0.99, 0.95, 0.9, 0.8, 0.7, 0.6, and 0.5.

For values of $\alpha < 1$, we executed FastSinkSource to convergence using our new test. Specifically, we used as many iterations as needed to fix the relative rankings of the subset of nodes that were either positive or negative examples in the left-out species. Note that we required knowledge only of whether or not a gene was in this subset, i.e., we did not know which gene was a positive and which was a negative example. For $\alpha = 1.0$, our new strategy for testing convergence is not applicable since we can only prove a trivial bound of unity between a node's final score and its score after $i$ iterations. Hence, we chose to stop FastSinkSource after 2,000 iterations. We selected this value since we observed that to reach a value of $\epsilon = 1 \times 10^{-4}$, FastSinkSource with $\alpha = 1.0$ required a median of 1,833 iterations, with a median absolute deviation (MAD) of 844.5.

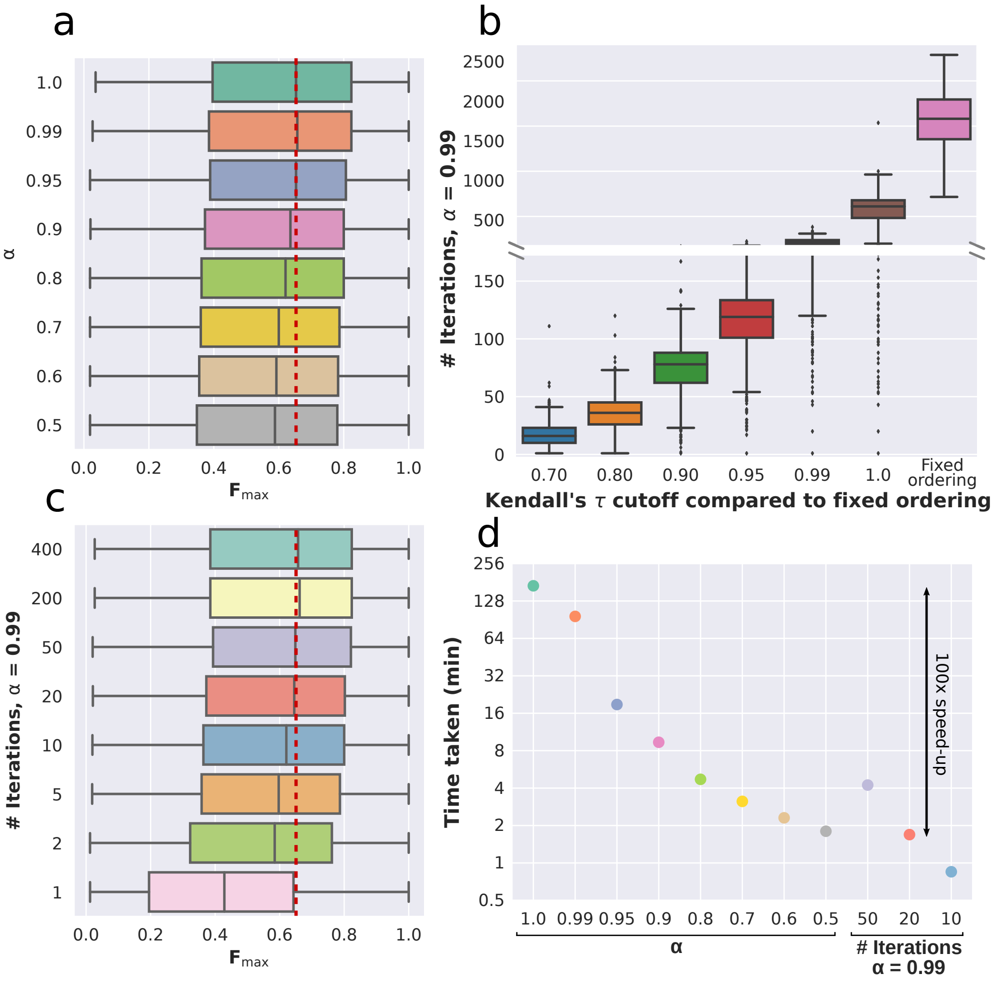


Figure S5: Trade-off between accuracy and speed of FastSinkSource for LOSO validation on the SSN-T+SSN-C+STRING-T+STRING-C (i.e., full) network. (**a**) Variation of $F_{\max}$ distributions with $\alpha$. The vertical red dotted line represents the median $F_{\max}$ for $\alpha = 1.0$. (**b**) Number of iterations required to fix the rankings of the left-out positive and negative examples, or to reach a specified value of Kendall's $\tau$ in comparison to the fixed ranking. (**c**) Variation of $F_{\max}$ distributions with the number of iterations ($\alpha = 0.99$). The vertical red dotted line represents the median $F_{\max}$ for $\alpha = 1.0$ in panel (a). (**d**) Total time taken by FastSinkSource while varying $\alpha$ ($F_{\max}$ values shown in (a)) or the number of iterations with $\alpha = 0.99$ ($F_{\max}$ values shown in (c)). Colors are the same as in (a) and (c).

We explored the effect of varying $\alpha$ on the accuracy of FastSinkSource by comparing the resulting distributions of $F_{\max}$ values over the BP GO terms we tested. The median $F_{\max}$ value did decrease gradually as we decreased $\alpha$ (Figure S5a). For $\alpha \geq 0.99$, we observed a statistically indistinguishable difference compared to $\alpha = 1.0$ (BF-corrected one-sided Wilcoxon signed-rank test $p$-value $> 0.25$). We chose to use $\alpha = 0.99$ for subsequent analyses.

Next, we considered varying the number of iterations for which we ran FastSinkSource. We first executed FastSinkSource to convergence, i.e., till the node orderings were fixed. We then re-executed FastSinkSource and for each $i > 0$, we compared how similar the node rankings after iteration $i$ were to the fixed ordering. We computed this similarity using Kendall's $\tau$, a measure of rank correlation. Given two different rankings of the nodes, this measure counts the fraction of node pairs that are not inverted, i.e., the fraction of node pairs $u$ and $v$ such that $u$ appears before $v$ in both rankings or vice-versa. Thus, the closer $\tau$ is to unity, the more similar are the two rankings. For each GO term, we recorded the number of iterations needed to reach a Kendall's $\tau$ value of 0.70, 0.80, 0.90, 0.95, 0.99, and 1.0 (Figure S5b). The median number of iterations required to fix the node ordering completely was 1455. Interestingly, we observed that FastSinkSource computed this ranking earlier with a median of 610 iterations (Figure S5b, Kendall's $\tau = 1.0$); subsequent iterations did not change the ordering but were necessary to guarantee that it would not be modified by more computations. Even more strikingly, after a median of 16 and 36 iterations, Kendall's $\tau$ was already as high as 0.70 and 0.80, respectively. These results suggested that while about 1500 iterations may be required to provably fix the rankings of the left-out positive and negative nodes, reducing the number of iterations by a factor of 40 does not have a major impact on the relative orderings of most of the positive and negative node pairs. Therefore, fewer iterations may not be deleterious to the accuracy of FastSinkSource during evaluation.

To test this hypothesis, we fixed $\alpha = 0.99$, ran FastSinkSource for eight different numbers of iterations (400, 200, 50, 20, 10, 5, 2, and 1), and recorded the $F_{\max}$ values after LOSO validation (Figure S5c). The $F_{\max}$ distribution for 400 iterations was statistically indistinguishable from the distribution for each of the next three largest number of iterations (200, 50, and 20, BF-corrected one-sided Wilcoxon signed-rank test $p$-value $> 0.25$). We concluded that decreasing the number of iterations by a factor of 20 (from 400 to 20) did not affect the overall distribution of LOSO performance. The running time improved by a factor of 100 (169 minutes for SinkSource with $\alpha = 1.0$ for 2,000 iterations vs. 1.69 minutes for FastSinkSource with $\alpha = 0.99$ for 20 iterations, Figure S5d). We chose to run FastSinkSource with $\alpha = 0.99$ and 20 iterations in all subsequent analyses for BP GO terms. For MF terms, we chose to keep the same parameter values for FastSinkSource.

#### S4.4 Parameter Selection for BirgRank

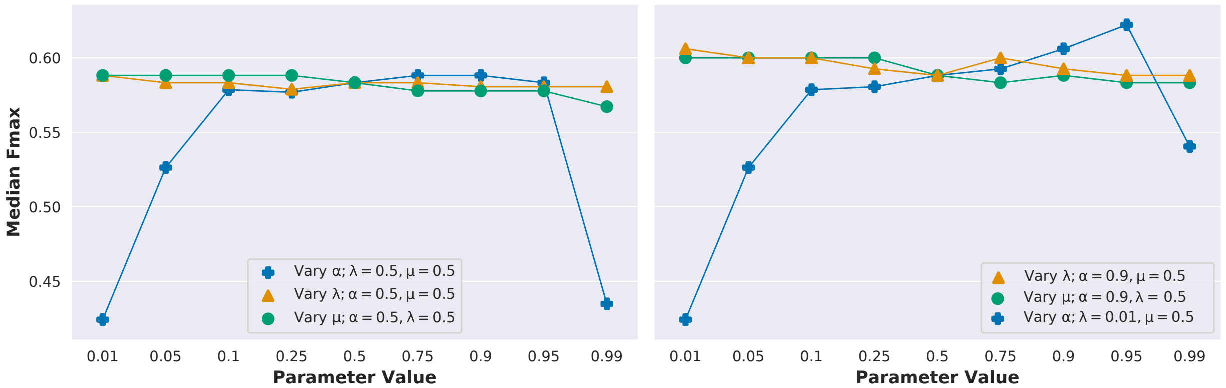


Figure S6: BirgRank parameter comparison of the $F_{\max}$ results of the EXPC LOSO evaluation on the full network. Related to Figure 3. (**a**) We first fixed every pair of parameters at 0.5 and then tested a range of values for the third parameter. (**b**) We then set $\lambda = 0.01, \mu = 0.5$ and varied $\alpha$, then set $\alpha = 0.95, \mu = 0.5$ and varied $\lambda$, and finally set $\alpha = 0.95, \lambda = 0.01$ and varied $\mu$.

To find the best parameters for BirgRank, we performed LOSO validation (BP EXPC annotations, SSN-T+SSN-C+STRING-C+STRING-T network) and varied the parameters $\alpha$, $\mu$, and $\lambda$, which control the RWR restart probability, proportion of the flow within $G$ vs. to $H$, and the direction of flow within the GO hierarchy, respectively. We did not need to vary the parameter $\theta$ (percentage of restart to a given node vs. its annotations) since LOSO evaluation removed all annotations from the nodes on which we ran BirgRank. We started by varying $\alpha$ with $\lambda = 0.5$ and $\mu = 0.5$, $\lambda$ with $\alpha = 0.5$ and $\mu = 0.5$, and $\mu$ with $\alpha = 0.5$ and $\lambda = 0.5$. We observed that $\alpha$ had the greatest affect on the median $F_{\max}$ (blue points in Figure S6a), whereas $\mu$ and $\lambda$ had a minor affect (green and yellow points in Figure S6a). Therefore, we selected the value of $\alpha = 0.9$ which resulted in the highest median $F_{\max}$ value for further analysis. With $\alpha = 0.9$, we set $\mu = 0.5$ and varied $\lambda$, and varied $\mu$ with $\lambda = 0.5$ (yellow and green points in Figure S6b, respectively). We observed the highest $F_{\max}$ of 0.61 with $\lambda = 0.01$. We selected this value of $\lambda$ and repeated our experiments with $\alpha$ to test our earlier settings for this parameter. Here we found that $\alpha = 0.95$ resulted in the highest median $F_{\max}$ of 0.64 (blue points in Figure S6b). Finally, we repeated the test for $\mu$ with the best values for the other parameters, $\lambda = 0.01$ and $\alpha = 0.95$, and observed a decrease in the median $F_{\max}$ for all parameter values except $\mu = 0.5$ (not shown). We chose to use the parameter values $\alpha = 0.95$, $\lambda = 0.01$, and $\mu = 0.5$ for all subsequent analyses.

### S5 Discussion of BirgRank Results

In Figure 3, many points are missing for BirgRank. As mentioned in the figure caption, this is due to our imposed limit of 72 hours of running time. For the first seven columns of Figure 3(a,d), we computed the scores of only the positive and negative examples of the left-out species. For the last two columns (i.e., SSN-Nbrs), we needed to compute scores for all of the genes in the core species for them to be transferred to a given left-out species, meaning we would have to compute scores for all genes of 24 species (25 core species minus one), repeated 24 times, which surpassed the time limit.

In Figure 3(c,f), the missing points (i.e., excessive running time) in the first seven columns were due to the large number of positive and negative examples in the 175 target species for which we needed to compute prediction scores. For the last two columns, however, the scores for the genes in the 25 core species only needed to be computed once, after which they could be transferred to the target genes using the weighted average.

Besides the running time, BirgRank did not perform better than the other propagation methods in our evaluations. One of the evaluations performed by Jiang *et al.* involved withholding all annotations from a given percentage of proteins, which is similar to our LOSO validation. They observed that in this evaluation, their methods (BirgRank and AptRank) were not able to outperform GeneMANIA and other propagation methods, which parallels our results.
